# Pea-Saving Partners: Bacillus and Pseudomonas combat downy mildew in pea crops

**DOI:** 10.1101/2025.01.04.631291

**Authors:** Emeka Chibuzor Okechukwu, Catherine Jimenez-Quiros, Ömür Baysal, Süreyya Kocamaz, Burhan Arıkan, Anne Webb, Thomas Wood, Sanu Arora, Claire Domoney, David J. Studholme, Mahmut Tör

**Author notes:** Corresponding author Address: Molecular Plant and Microbial Biosciences Research Unit School of Science and the Environment, University of Worcester, Worcester WR2 6AJ United Kingdom.

## Abstract

Downy mildew (DM) is a destructive disease that significantly reduces the yield and quality of important pulses (legumes) and horticultural crops, particularly during humid and cool seasons. This disease is caused by obligate and host-specific oomycete pathogens. Controlling the pathogen is challenging due to its long-term survival as spores and its rapid mutation. Use of chemical pesticides has been the most effective method to control DM pathogens, but their environmental hazards are a global concern. Current research is focused on exploring the potential of microbial biological control agents (MBCA), particularly rhizobacteria strains of the genera *Bacillus* and *Pseudomonas*, which have shown suppression of plant pathogens. However, to date, no MBCA has been reported to be effective against DM pathogens in pulses. We investigated the effectiveness of *Bacillus* and *Pseudomonas* strains as potential biopesticides against the pea downy mildew pathogen *Peronospora viciae* f. sp. *pisi* (*Pvp*). In our study, *in vitro* bioassays showed 100% inhibition of *Pvp* spore germination compared to the control. *In planta* antagonism assays further demonstrated significant suppression (>80%) of *Pvp* sporulation in pea plants sprayed with strains of *Bacillus velezensis* or *P. fluorescens* or their filtrates. The drench application also showed significant effects where either a *Pseudomonas* or cold-adapted *Bacillus* strain was used. We observed a synergistic effect for the dual foliar application of the microbes compared to individual application (27.6 to 46.7% suppression). Furthermore, the results from the molecular biomass analysis were consistent with the results of the sporulation assays. This demonstrates the strong interactive and promotive benefits of using *Bacillus* and *Pseudomonas* as biocontrol agents Based on these results, we conclude that these MBCAs could be effective in combatting *Pvp* infections in the field.

## Introduction

Plant pathogens have been serious and persistent threats to global crop yield and quality (Ristaino et al., 2021; Yang et al., 2022). Along with pests, they cause up to 40% crop loss globally each year, which cost the global economy billions of dollars (Jamiołkowska, 2020; Pandit et al., 2022). The global concern is not only for the present threats from the existing plant pathogens that have persisted for centuries, but also from emerging ones as occasioned by climate change (Burdon & Zhan, 2020; Corredor-Moreno & Saunders, 2020; Ristaino et al., 2021; Velasquez et al., 2018). Pathogenic attacks are one of the primary causes of global food insecurity, and their impacts could worsen by 2050 when the world’s human population is projected to reach approximately 10 billion. (Velasquez et al., 2018; Zhao et al., 2022). This further highlights the global urgency of reducing pathogen-induced yield loss (McDonald & Stukenbrock, 2016; Savary et al., 2019).

Downy mildew (DM) is one of the world’s most devastating plant diseases; it seriously reduces yield (up to 80%) and quality of globally important pulses, vegetables, fruits, and ornamentals, most notably during humid-cool seasons that are usually synchronised with the cropping seasons (Salcedo et al., 2021; Siddaiah et al., 2017). The disease is caused by obligate biotrophic pathogens that exhibit host-specificity (Choi & Thines, 2015; Thines, 2009; van Damme et al., 2009). Some of the common downy mildew pathogens are *Peronospora viciae* f. sp. *pisi* (pea), *P. viciae* f. sp. *fabae* (faba beans), *Hyaloperonospora brassicae (brassica), H. parasitica (Capsella bursa-pastoris), P. belbahrii (basil), P. destructor* (onion), *P. manshurica* (soyabean), *P. effusa* (spinach), *Bremia lactucae* (lettuce) and *Plasmopara viticola* (grapevine), *Pseudoperonospora cubensis* (cucurbits) and *Plasmopara halstedii* (sunflower) (Salcedo et al., 2021; Thines, 2009; Tor et al., 2023). They attack above-ground plant parts such as the leaves, stems, flowers, pods, and fruits (Koledenkova et al., 2022). The effects on plants include stunted growth, distortion and discoloration of leaves, and typical fluffy mold-like growth on the surface of the leaves (Bandamaravuri et al., 2020). The pathogens are resilient and adaptable to new environments (Delmas et al., 2016), due to their ability to survive as long lasting spores (oospores) under harsh conditions or in absence of host plants and to rapidly mutate to evade or overcome pesticides or host defences (Koledenkova et al., 2022).

For many years, chemical pesticides such as Wakil XL (metalaxyl-M, fludioxonil and cymoxanil) have been the most effective method to control DM pathogens such as *P. viciae* f. sp. *pisi* (*Pvp*) in peas. However, indiscriminate and continuous use of these chemicals has caused a lot of short and long-term hazards particularly to the environment and ecosystem, and accumulation of their associated toxic residues in the food chains pose serious threats to human and animal health, and wellbeing (Aktar et al., 2009; Damalas & Eleftherohorinos, 2011; Lahlali et al., 2022). Strict regulations have been implemented on the timing and usage of pesticides in different countries and more restrictions will follow with a long-term aim of achieving full-scale global sustainable crop production (Lahlali et al., 2022). Towards this aim, research is increasingly focusing on developing new alternatives for managing plant pathogens that will not only be effective, but also safe, sustainable, and eco-friendly. Some of the non-chemical pesticides that hold great promise are biological/biocontrol agents or their byproducts (Jimenez-Quiros et al., 2022; Pandit et al., 2022), plant extracts (Cowan, 1999), phage therapy (Erdrich et al., 2024; Villalpando-Aguilar et al., 2022) and more recently small interfering RNAs, popularly called spray-induced gene silencing (Bilir et al., 2022).

Microbial biological control agents (MBCA) have been the most broadly studied and utilized biopesticides (Jaiswal et al., 2022). Among them, rhizobacteria of the genera *Bacillus* and *Pseudomonas* have been shown to suppress a wide range of plant pathogens of different phlya/kingdoms (Dragana et al., 2017; Gao et al., 2012; Mnif & Ghribi, 2015). We previously demonstrated that a strain of *Bacillus velezensis* (EU07), whose genome was sequenced (Baysal et al., 2024), effectively controlled *Fusarium graminearum,* the pathogen that causes *Fusarium* head blight disease in cereals (Jimenez-Quiros et al., 2022). Although some non-pathogenic *Fusarium* and *Trichoderma* isolates have been reported to be effective against some downy mildew pathogens (Bakshi et al., 2001; Nandini et al., 2021; Núñez-Palenius et al., 2022), there are no reports of biocontrol of a downy mildew pathogen that affects important legume crops such as peas. To address this critical research gap, this study aimed to investigate the effectiveness of *Bacillus* and *Pseudomonas* strains as potential biopesticides against *Pvp*.

## Materials and Methods

### Biological agents used and preparation of inoculum

We tested two *Bacillus velezensis* strains that are commercially available as biocontrol products: Serenade (QST713) and TAEGRO370 (FZB24). Strain FZB24 is the type strain of *B. amyloliquefaciens* subsp. *plantarum* (Borriss et al., 2011) but this taxon is now properly considered as belonging to the species *B. velezensis* (Parte, 2018). We also tested a non-commercial strain, *B. velezensis* EU07 (Baysal et al., 2013; Jimenez-Quiros et al., 2022 (*Bacillus velezensis*), whose genome sequence is almost identical to that of QST713 (Baysal et al., 2024). We also evaluated the cold-adapted *Bacillus* strains K7, K9, K11, K12 and B2-6 isolated from persimmon (tree) leaf litter in Tarsus, Mersin, Turkey (at an altitude of 1200 m) during the cold season after the snow melted, and *Pseudomonas fluorescens* strain LZB 065, procured from Blades Biological Ltd, UK. The bacteria were streaked on Luria-Bertani (LB) agar (Bertani, 1951) and incubated at 15°C or 28°C for 2 days to produce single colonies. After genetic identity verification through PCR and sequencing procedures, a colony from each strain (after genetic identity verification through PCR and sequencing procedures) was used to produce glycerol stocks that were flash-frozen in liquid nitrogen and stored at −80°C until needed. To make bacterial broths, bacteria were streaked on LB plates from glycerol stocks and a single colony was grown in liquid LB media in a shaker (15°C or 28°C, 220 rpm) OD_600_∼2 was obtained.

### Re-verification of *Bacillus* QST713, FZB24 and EU07 Strains

We re-confirmed the genetic identity of the *Bacillus* QST713, FZB24 and EU07 strains through a combination of colony PCR and Sanger sequencing techniques. Specifically, the PCR protocol for amplification of 16sRNA genes involved an initial denaturation step at 94°C for 3 minutes, followed by 35 cycles of denaturation (94°C for 30 seconds), annealing (55°C for 30 seconds), and extension (72°C for 1 minute) and final extension (72°C for 5 minutes). Gel electrophoresis validated the expected band size of the PCR products. Subsequently, the purified bands underwent Sanger sequencing. To further validate our findings, we performed BLASTN analysis against known *Bacillus* 16S rRNA gene sequences in the NCBI databases (Altschul et al., 1990; Baysal et al., 2024; Jimenez-Quiros et al., 2022). Finally, these re-verified bacterial colonies were maintained as glycerol stocks, and used to generate bacterial broth used for further steps of the studies. The primers used are presented in Supplemental Table S1.

### Whole-genome sequencing of *Pseudomonas* and cold-adapted *Bacillus* Strains

Our initial screening of cold-adapted *Bacillus* strains showed K11 was the best out of five cold-adapted strains tested, and therefore we concentrated on this. We carried out whole genome sequencing of the *Bacillus* K11 and *P. fluorescens* LZB 065 strains as they have not been sequenced prior to this study. Overnight liquid cultures of the bacteria were produced from their single colonies. To harvest the pellets, 2ml culture was centrifuged for 5 minutes at 8,000 x g and the supernatant was discarded. Genomic (g) DNA was extracted following the steps explained in Meridian Bioscience ISOLATE II Genomic DNA Kit (Scientific Laboratory Supplies Ltd, UK). The quality of the gDNAs was assessed using the Agilent 4200 TapeStation to confirm that they meet the required standards for genomic sequencing. The samples were sent to Novogene, UK for whole-genome sequencing, generating150-bp paired reads via the Illumina NovaSeq 6000 instrument.

### Genome assembly and annotation

Prior to genome assembly, we filtered and trimmed the raw Illumina sequence reads using TrimGalore version 0.6.7, which incorporates Cutadapt version 3.5 (Krueger, 2019). The -q parameter was set to 30 and we used the --paired option. For *de-novo* assembly of these processed reads, we used Unicycler v. 0.5.1 (Wick et al., 2017), which incorporates SPAdes v. 4.0.0 (Bankevich et al., 2012). The command line was: “unicycler -1 short_reads_1.fastq.gz -2 short_reads_2.fastq.gz -o output_dir”. We submitted the resulting genome sequence assemblies to GenBank (Benson, 2004) via the NCBI Submission portal (Sayers eet al, 2019). Genome annotation was generated by the NCBI’s PGAP pipeline v. 6.8 (Tatusova et al., 2019).

### Maintenance and propagation of *Pvp* on pea plants

The purified *Pvp* isolate 20-1-3 (DM3)’, originally collected in 2020 from infected pea plants in Cambridge, UK, was obtained from the culture collection of NIAB and used throughout this study. The shoots were harvested from *Pvp*-infected plant of pea cultivar (cv.) Maro were harvested and placed in a beaker with sterile water. The beaker was gently agitated to shake spores off the shoots. The spore suspension was then filtered through a layer of Miracloth into clean glassware. The spore count was determined using a haemocytometer under a light microscope and adjusted to the required concentration. The spore solution obtained was used to inoculate 4-day old pre-germinated pea seeds. The seedlings were immersed in the spore solution for 30 minutes with gentle shaking every 5 minutes to ensure uniform inoculation. They were then immediately sown in standard compost (Levington Advance Seed & Modular F2S Compost - Plus Sand) and transferred to a growth cabinet (16°C, 12 hours light and 12 hours dark). Ten days post inoculation (dpi), the inoculated plants were covered with transparent lids (with the edges sealed with electric tape) for 2 days to aid the sporulation of the pathogen. The spores formed were harvested and either used for experiments or re-propagated to maintain the pathogen on the host as summarised in Fig. 1.

**Figure 1.**
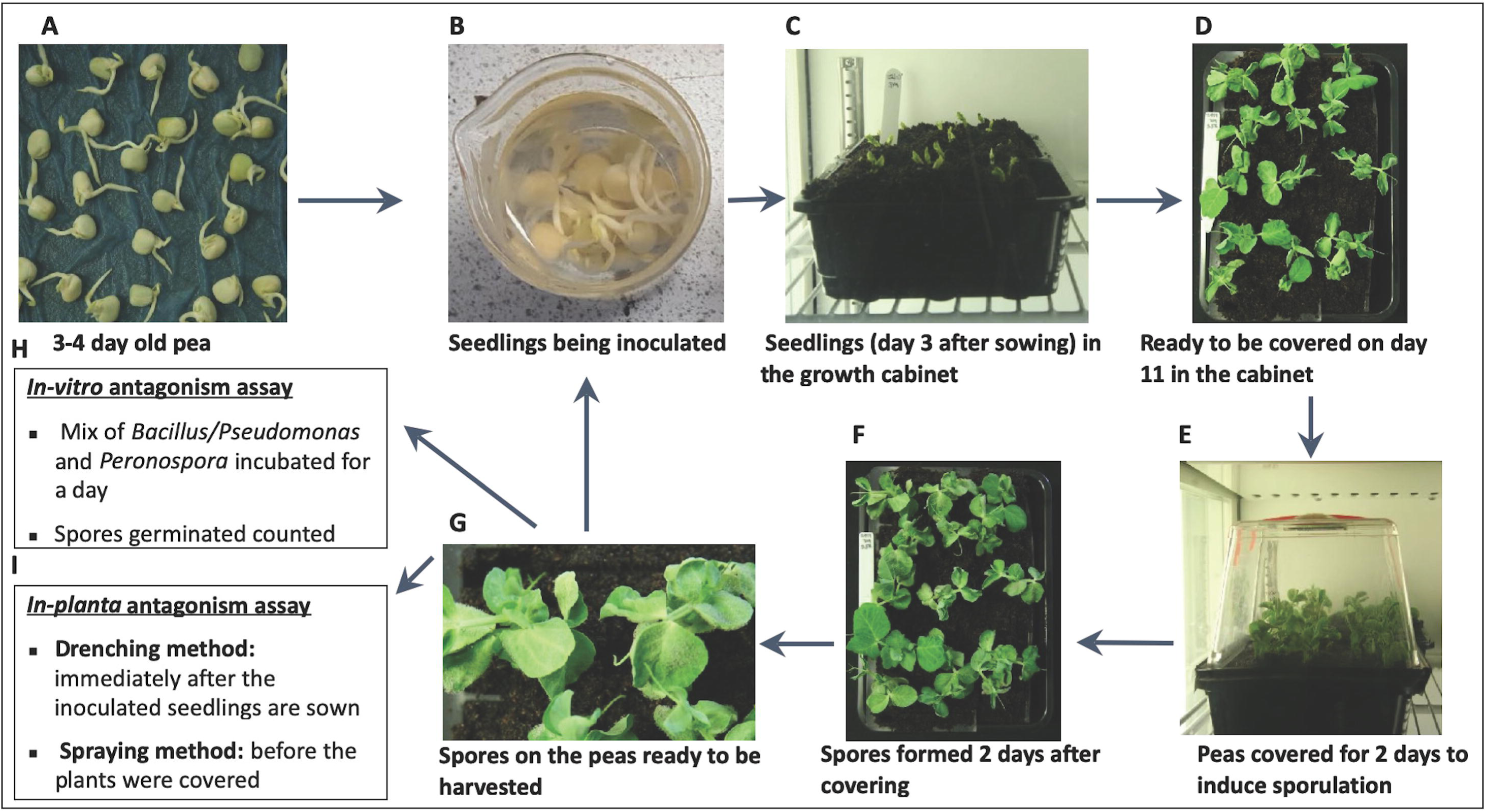
I**noculation of pea seedlings with *Pvp*. A)** Germinated pea seedlings ready for the *Pvp* inoculation. **B)** Seedlings were treated with *Pvp* spores (25,000 spores/ml) for 30 minutes for inoculations. **C)** *Pvp*-inoculated seedlings growing in the growth cabinet. **D**) Inoculated plants ready to be covered. **E)** Plants covered with a transparent lid to maintain humidity and induce sporulation of *Pvp*. **F-G**) Sporulation occurred after covering the trays. **H-I)** Spores were harvested and used for the biocontrol antagonism assays.

### *In vitro* antagonism bioassays of *Bacillus* and *Pseudomonas* strains on *Pvp* spore germination

The *Pvp* spores were harvested and cleaned by centrifugation at approximately 3000 rpm for 3 minutes, and washing in ice-cold water, repeated twice, followed by resuspension in water. Full-strength cultures (OD_600_ ∼2) of the biocontrol *Bacillus* strains were centrifuged at 14,000 rpm for 5 minutes to separate the cells (pellets) from the filtrates. Different concentrations of the filtrates (100%, 50%, 25%, and 12.5%) and bacterial cells (OD_600_ of 1, 0.5, and 0.25) were separately mixed with the *Pvp* spores (final concentration of 25,000 spores/ml). One hundred µl from each mix was plated on a microscope slide (two spots per slide) placed on a transparent petri dish. The lids were covered, and the Petri dishes were placed in the growth cabinet (16°C, 12 hours light and 12 hours dark) for a day to allow spore germination. To quantify the antagonistic effect of Bacillus/Pseudomonas on *Pvp*, the percentage of germination from both treated and untreated spores were measured and compared using statistical analysis.

### *In planta* antagonism assay of *Bacillus* and *Pseudomonas* strains on *Pvp* development

Two different methods were used: drenching and foliar spray applications. For the drenching method, 4-day old pea seedlings were inoculated with *Pvp* spores (25,000 spores/ml) and planted in 15 multi-cell trays filled with standard compost (Levington Advance Seed & Modular F2S Compost - Plus Sand). Each seedling was drenched with 25ml of biocontrol full-strength culture (OD_600_ ∼ 2) or LB medium as a control. The plants were then moved to a growth cabinet (16°C, 12h light and 12h dark). Ten days after inoculation, the trays of plants were covered with transparent lids for 2 days to allow the pathogen to sporulate. The inoculated plants were then harvested, and spores were counted.

For the foliar spraying assay, the filtrates (supernatant after centrifugation and filtering cultures through 0.22µm filters (EMD Millipore Millex™) and cells (pellets resuspended in water) were sprayed on 10-day-old *Pvp*-inoculated pea plants using an electric atomizer. Each plant was sprayed with 20ml (supplemented with 0.05% silwet L-77) of the biocontrol cells or their filtrates. Full strength filtrates and the cells with OD_600_ 5 were used. LB and water were applied as controls for biocontrol filtrates and cells, respectively. The plants were allowed to air-dry for 5 minutes, covered with lids, and moved back to the cabinet for a further 2 more days to allow the pathogen to sporulate. The sporulated plants were then harvested, and spore counts were carried out.

### DNA Extraction

DNA was extracted from *Pvp*-inoculated pea plants that were either drenched or sprayed with the biocontrol treatment or experimental control (LB or water). The extraction was performed using the traditional cetyl trimethylammonium bromide (CTAB) method with polyvinylpyrrolidone (PVP), as described by Koh et al. (2021).

### Biomass analysis using quantitative PCR

The quantitative PCR (qPCR) technique was used to measure the *Pvp* mycelial biomass using qPCRBIO SyGreen Mix Lo-ROX (PCR Biosystem, UK) as the preferred master mix. For each sample, a reaction mix of 20 μl prepared, which included 10 μl of SyGreen Mix Lo-ROX, 0.8 μl of 10 μM forward primer, 0.8 μl of 10 μM reverse primer, 1 μl of 100ng DNA template, and 7.4 μl of nuclease-free H_2_O was prepared. The PCR reaction was performed in a Roche 480 II thermocycler with the following program: 3 minutes at 95°C for polymerase activation, followed by 40 cycles of (5 sec at 95°C, 20 sec with a touchdown step size of 0.8°C from 65°C to 60°C) for denaturation and annealing/extension, and 1 min cooling down at 40°C. The *Pvp-Actin* primer pair was used to amplify a unique region of the *Pvp-Actin* gene, and the *Ps-Actin* (pea-Actin) primer pair was used for normalization (housekeeping) of host DNA. Three biological replicates, each with two technical repeats, were used. To compare the relative abundance of *Pvp-Actin* to *Ps-Actin* for the biocontrol treated and the mock treated samples, the fold change was calculated relative to the control (2^-ΔΔCT) as explained by Schmittgen and Livak (2008).

### Statistical Analysis

A two-tailed, unpaired, heteroscedastic t-test was used to determine if there was a significant difference between the biocontrol and experimental control. The means and standard errors were displayed in plots. Bar plots were generated using Microsoft Excel version Version 16.89.1(24091630), while R software version 4.4.1 (Race for Your Life), RStudio IDE version 2024.04.2+764 (2024.04.2+764) and ggplot (Wickham, 2016) were used to construct the box plots.

### Bioinformatics

For identification of bacteria to species level, we uploaded genome assemblies to the Type Strain Genome Server (TYGS) (Meier-Kolthoff et al., 2019; Meier-Kolthoff et al., 2022) at https://tygs.dsmz.de/user_requests/new.

To calculate average nucleotide identities, we used FastANI version 1.33 (Jain et al., 2018). To generate a maximum-likelihood phylogenetic tree based on genome-wide single-nucleotide variants we used PhaME version 1.0.2 (Shakya et al., 2020) with FastTree version 2.1.11 (Price et al., 2010). This generated a tree, which we graphically rendered using the Interactive Tree of Life (iTOL) 7.0 (Letunic and Bork, 2021). Essentially, we used the same protocols for ANI and phylogenomics analysis as described previously (Baysal et al., 2024) but included additional genome sequences with strain K11.

### Accession numbers

All genome sequence data have been deposited in public databases under the BioProject accession PRJNA1150624. Raw sequence reads are deposited in the Sequence Read Archive (Kodoma et al., 2012) under the following accession numbers: SRX25802839 (QST713), SRX25802838 (FZB24), SRX25793480 (K-11) and SRX25793481 (LZB 065). Annotated genome assemblies are deposited in GenBank under the accession numbers GCA_045108535.1 (QST713), GCA_045108515.1 (FZB24), GCA_041520185.1 (K-11), GCA_041521055.1 (LZB 065).

## Results

### *Bacillus and Pseudomonas* strains inhibit germination of *Pvp* spores *in vitro*

*Pvp* spores were grown on pea plants, harvested, and examined using *in vitro* antagonism bioassays. Three strains of *B. velezensis* (EU07, FZB24, QST713) and *P. fluorescens* were mixed with the *Pvp* spores and incubated on microscope slides overnight. The biocontrol agents completely suppressed *Pvp* spore germination (100%) when treated with bacterial cells at OD_600_ of 1 and 0.5. However, a lower bacterial cell concentration (OD_600_ of 0.25) did not show inhibitory effects. Similarly, filtrates of the *Bacillus/Pseudomonas* strains at 100% and 50% concentrations showed 100% inhibitory effects (Figs. 2 and 3, respectively). At the lowest concentration of 12.5%, there was still a significant reduction in spore germination percentage for *Bacillus* / *Pseudomonas* treated spores compared the control (LB medium), although some spores (2.5% to 29.5% relative to the control) were able to germinate.

**Figure 2:**
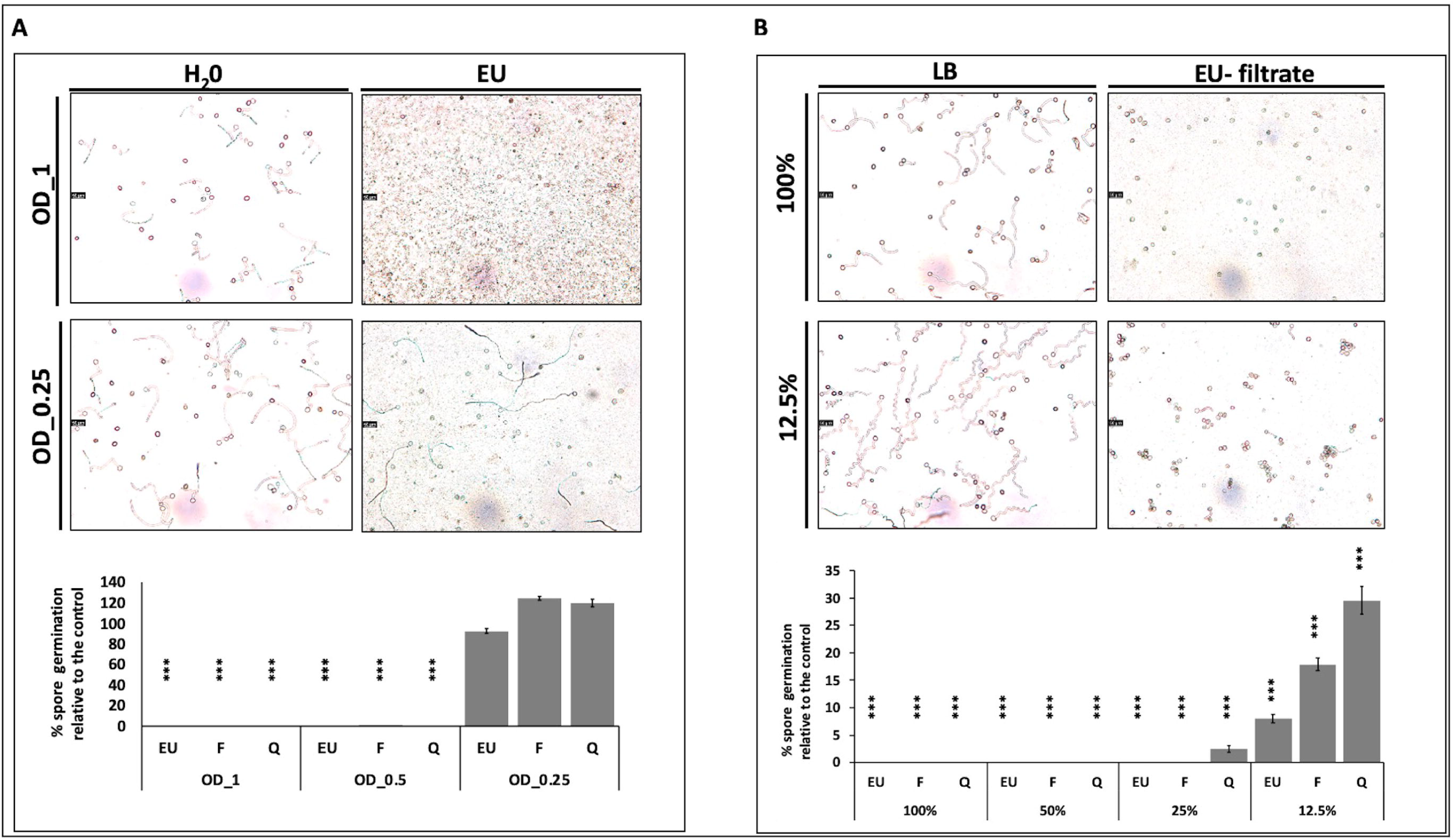
**Inhibitory effect of *Bacillus* strains o*n Pvp* spore germination**. *Pseudomonas* and the *Bacillus* strains EU, F, Q were tested. The effects of the biocontrol were presented relative to 100 % of the mock treatment. Antagonism assay of three *Bacillus* strains: cells (**A**) and filtrates (**B**) of varying concentrations on *Pvp* spore germination percentage. Water was used as the mock treatment for the bacterial cells, while LB was for the filtrates. Full-strength broths of the strains were centrifuged to separate the cells (pellets) from their filtrates. The filtrates concentrations tested were 100, 50, 25 and 12.5%, while the optical density (OD_600_) of bacterial cells examined were 1, 0.5 and 0.25. Mixtures of *Pvp* spore solution and the biocontrol were placed on a microscope slide in Petri dish and incubated overnight in growth cabinet. Percentage of germinated spores were calculated. The bar plots on the bottom of each panel showed data from one of the 3 independent biological repetitions. Error bars represent standard error from 3 technical replicates. T-test was used to compare the means for significant differences. ***=significant at p-value of <0.001.

**Figure 3:**
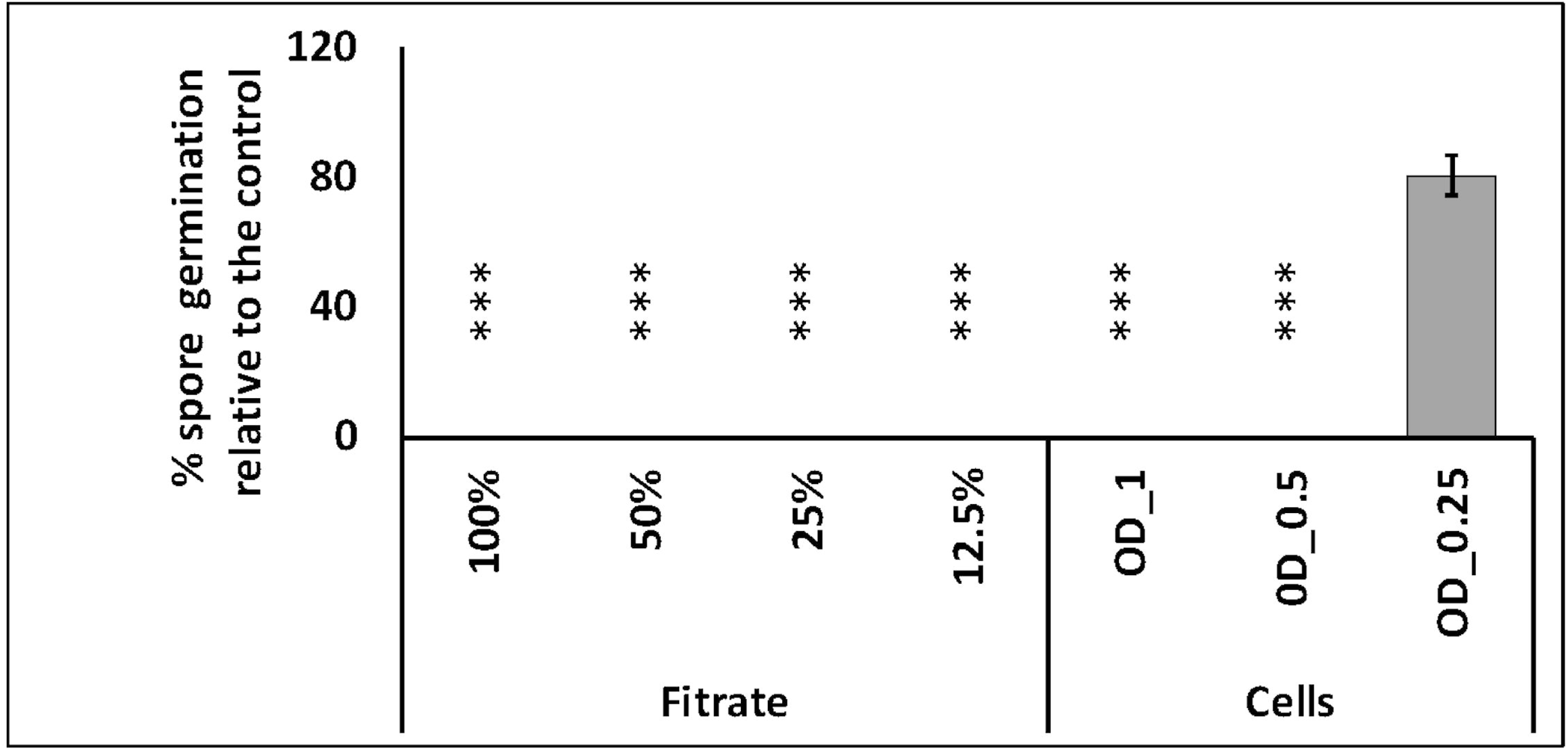
Inhibitory effect of *Pseudomonas* strain on *Pvp* spore germination. The effect of the biocontrol was presented relative to 100 % of the mock treatment. Antagonism assay of the biocontrol cells and filtrates of varying concentrations on *Pvp* spore germination percentage was tested. Water, the mock treatment for the bacterial cells; LB, for the filtrates was used. Full-strength broths of the strains were centrifuged to separate the cells (pellets) from their filtrates. The filtrates concentrations tested were 100, 50, 25 and 12.5%, while the optical density (OD_600_) of bacterial cells examined were 1, 0.5 and 0.25. Mixtures of Pvp spore solution and the biocontrol were plated on Petri dish and incubated overnight in growth cabinet. Percentage of germinated spores were calculated. The bar plots on the bottom of each panel showed data from one of the three independent biological repetitions. Error bars represent standard error from 3 technical replicates. T-test was used to compare the means for significant differences. ***=significant at p-value of <0.001.

### The pesticide Wakil XL coated pea seeds controls downy mildew

The most effective method for controlling downy mildew (DM) pathogen in pea crops has been through seed treatment with the pesticide Wakil XL, which contains metalaxyl-M, fludioxonil, and cymoxanil. In this research, Wakil XL was used as a positive control for the DM pathogen. Pea seeds coated with Wakil XL were pre-germinated, and the resulting seedlings were inoculated with the *Pvp* pathogen. As anticipated, the pea plants did not exhibit any symptoms of DM disease compared to the control plants, even when the *Pvp* spore concentration was doubled to 50,000 spores/ml (Fig. 4A-C). In contrast, plants from untreated seeds showed full pathogen sporulations and dissease symptoms (Figure 4D).

**Figure 4:**
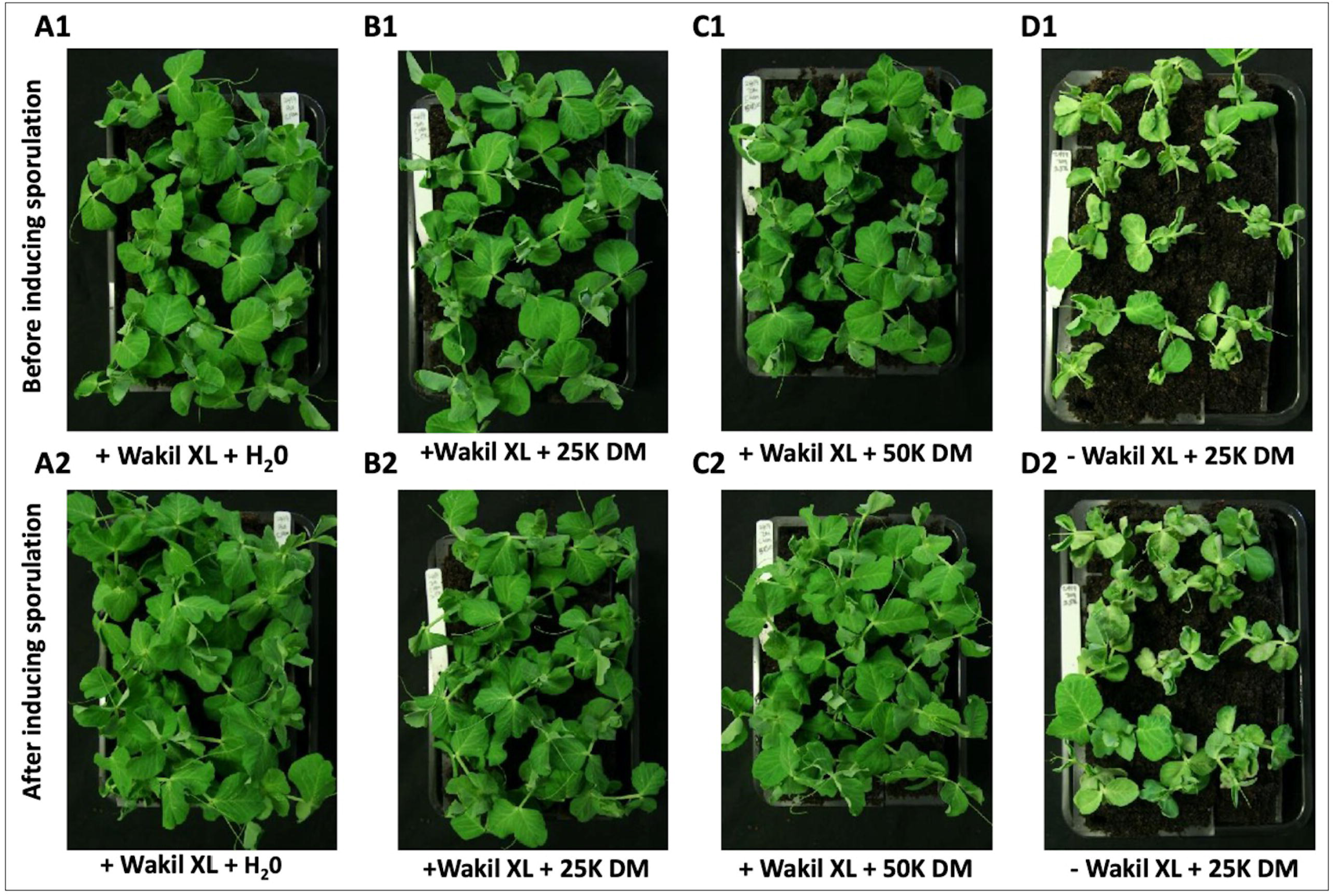
**Wakil XL was effective against *Pvp***. **A1)** Pea plants from the Wakil XL- coated seeds without *Pvp* inoculation, **B1)** pea plants from the Wakil XL-coated seeds with *Pvp* inoculation (25,000 spores/ml), **C1)** pea plants from the Wakil XL- coated seeds with *Pvp* inoculation (50,000 spores/ml), **D1)** pea plants from control seeds with *Pvp* inoculation (25,000 spores/ml), **A2)** pea plants from the Wakil XL- coated seeds without *Pvp* inoculation, **B2)** pea plants from the Wakil XL-coated seeds with *Pvp* inoculation (25,000 spores/ml), **C2)**, pea plants from the Wakil XL- coated seeds with *Pvp* inoculatin (50,000 spores/ml), and **D2)** pea plants from control seeds with *Pvp* inocultation (25,000 spores/ml). Images displayed in the top panel were taken 10 dpi, and pictures displayed in the lower panel were taken 2 days after covering the pea plants.

### Drenching the soil with *Bacillus and Pseudomonas* strains suppresses *Pvp* growth

We investigated whether using a biocontrol broth to drench the soil could inhibit the growth of *Pvp*. Spore count data showed that none of the three tropical *Bacillus* strains (EU07, FZB24, QST713) consistently or significantly reduced *Pvp* spore counts (Fig. 5). However, cold-adapted *Bacillus* and *Pseudomonas* strains significantly (p < 0.05) reduced pathogen sporulation by approximately 90% compared to the controls in three replicate experiments (Figs. 6 and 7, Supplemental Fig. S1).

**Figure 5:**
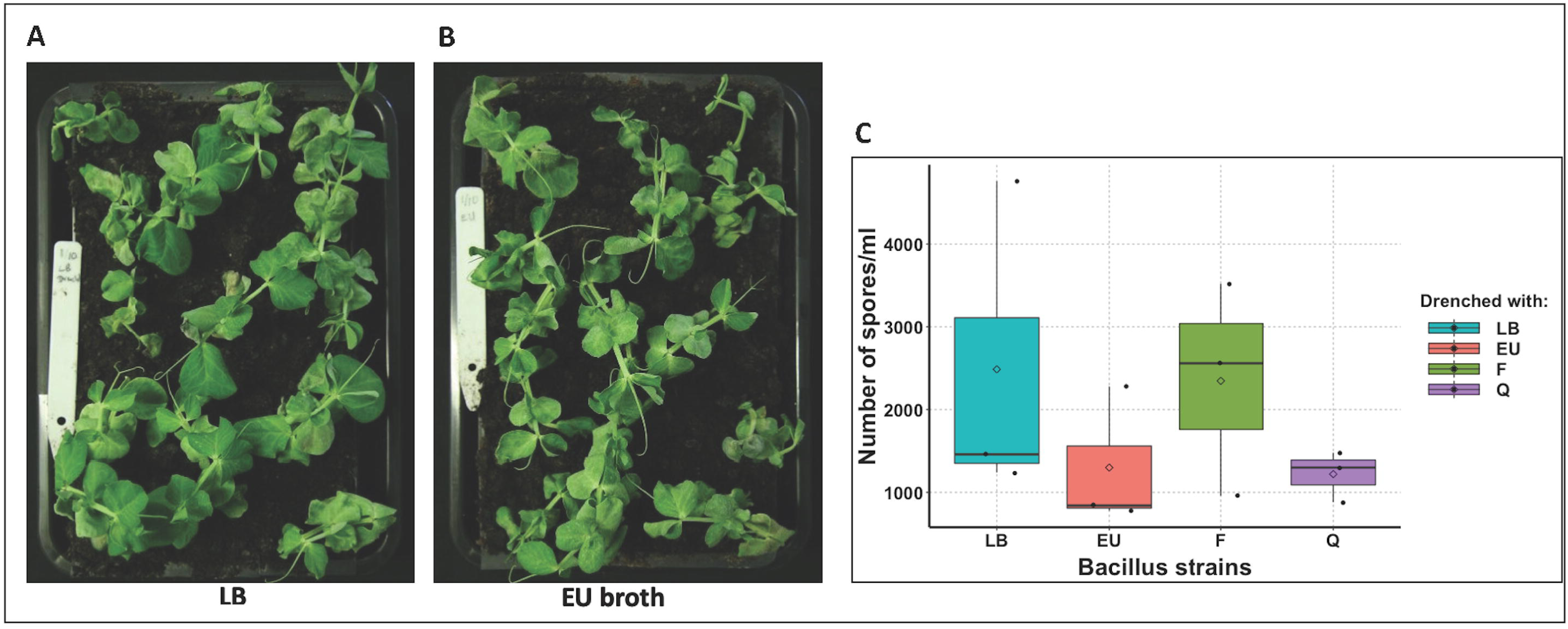
Antagonism assay of drench- application of tropical *Bacillus* broth on *Pvp*- inoculated pea plants. 4-day old pea seedlings were inoculated with *Pvp* spores and sown in a standard compost. Biocontrol broths or LB were applied immediately upon sowing the seedlings. After 10 days, the plants were covered for 2 days to induce *Pvp* spore formation. After the sporulation, plants drenched with LB (**A**) and EU- *Bacillus* broth (**B**) were photographed; mean spore counts for plants drenched with the three *Bacillus* strains (EU, F, & Q) and LB (control) are shown in box plots (**C**). Plots show data from one of the three independent biological repetitions. T-test was used to compare the means for significant differences.

**Figure 6:**
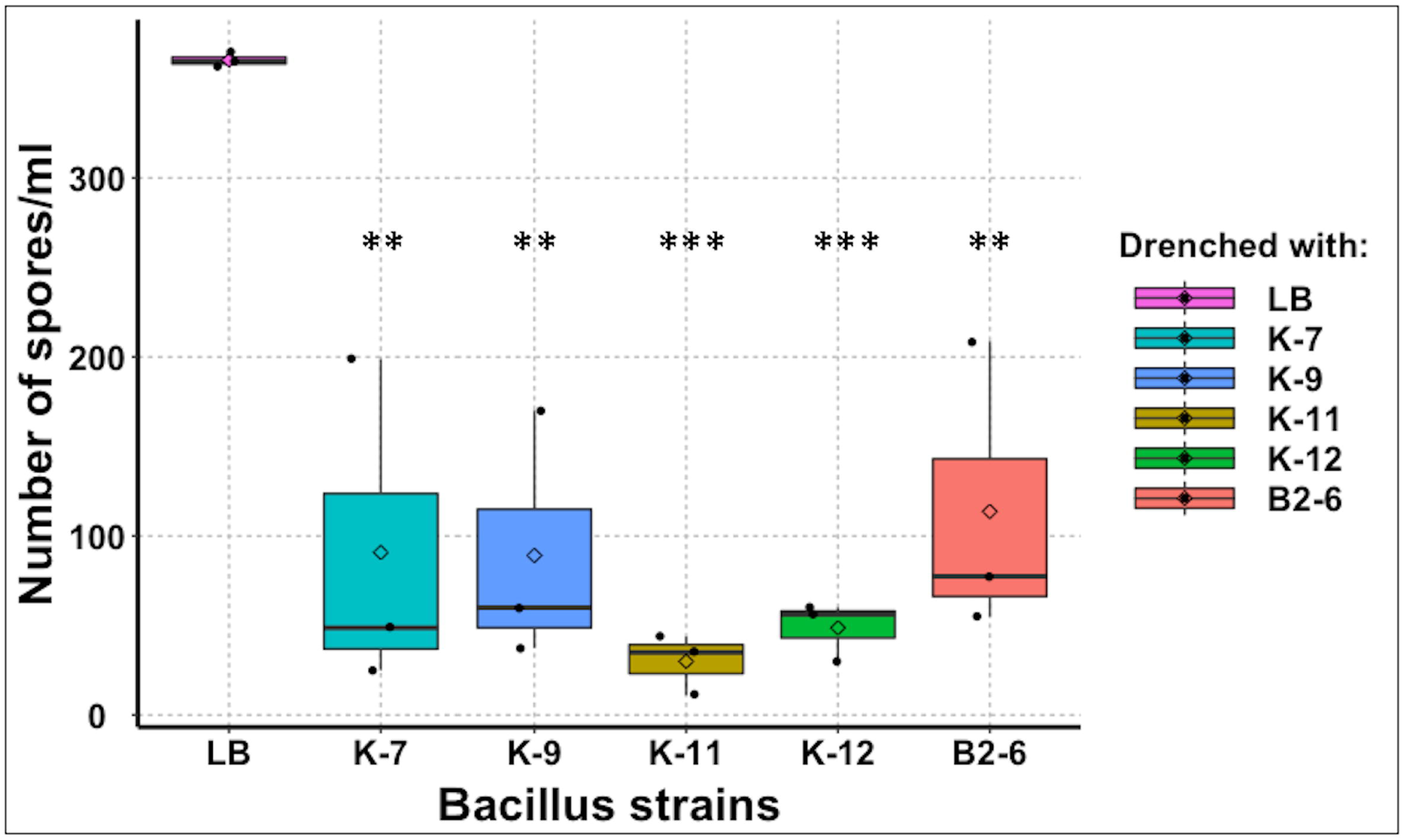
**Antagonism assay of drench- application of cold-loving *Bacillus* broth on *Pvp*- inoculated pea plants**. 4-day old pea seedlings were inoculated with *Pvp* spores and sown in a standard compost. Biocontrol broths or LB were applied immediately upon sowing the seedlings. After 10 days, the plants were covered for 2 days to induce *Pvp* spore formation. Mean spore counts for the plants drenched with the five *Bacillus* strains (K-7, K-9, K-11, K-12, & B2-6) and LB (control) are shown in box plots. Plots show data from one of the three independent biological repetitions. T-test was used to compare the means for significant differences.

**Figure 7:**
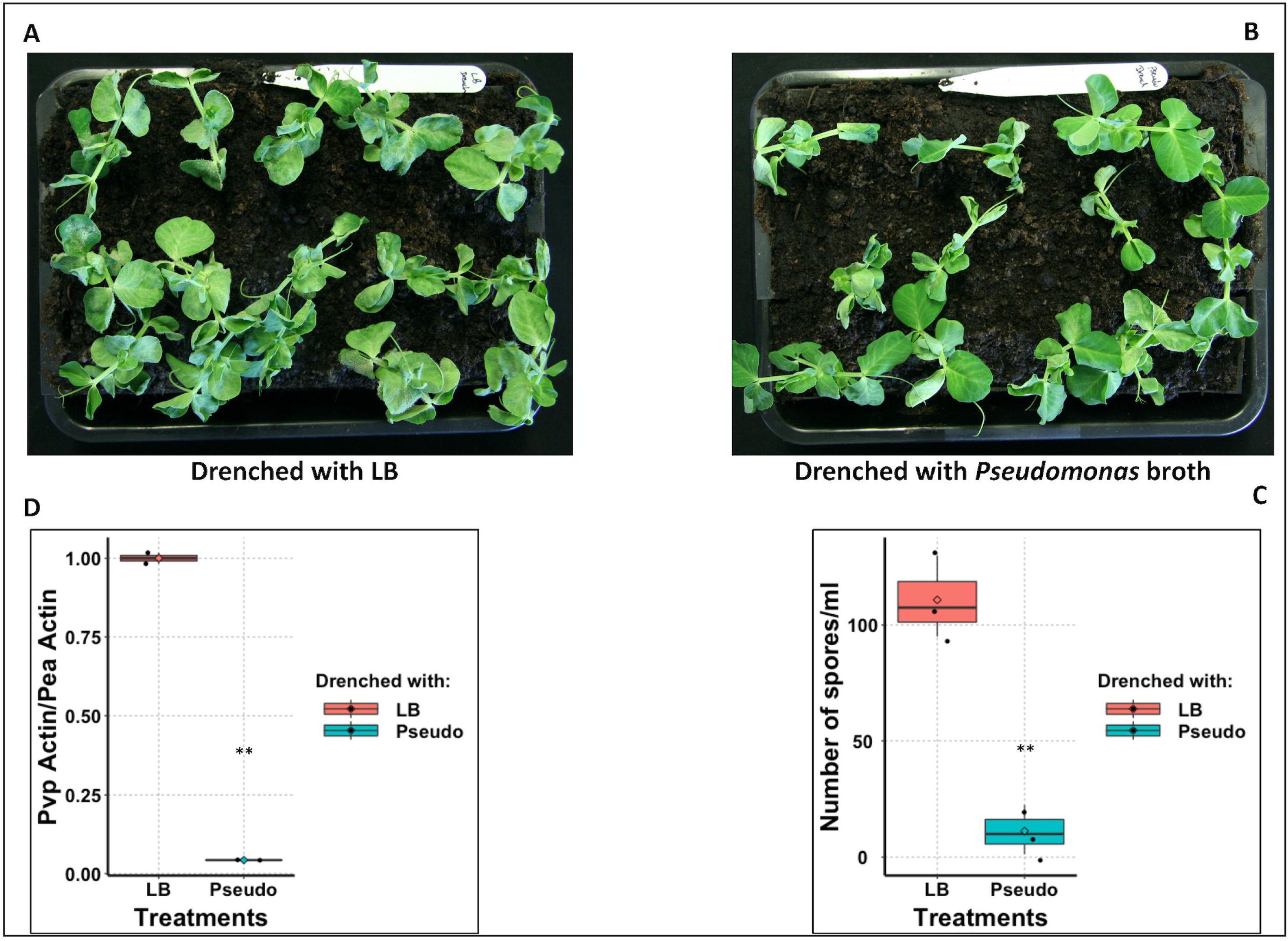
**Antagonism assay of drench-application of *Pseudomonas* broth on *Pvp*-inoculated pea plants**. 4-day old pea seedlings were inoculated with *Pvp* spores and sown in a standard compost. *Pseudomonas* broth or LB was applied immediately upon sowing the seedlings. After 10 days, the plants were covered for 2 days to induce *Pvp* sporulation. After the sporulation, plants drenched with LB (**A**) and *Pseudomonas* broth (**B**) were photographed; mean spore counts for plants drenched with *Pseudomonas* and LB (control) are shown in box plots (**C**). **D**: *Pvp* molecular biomass quantification in *Pseudomonas* and LB-drenched pea plants. *Pvp-Actin* primer pair was used to amplify a unique region of *Pvp* Actin, while a *Pea- Actin* primer pair was used for normalization (‘housekeeping’ control). The fold change of the *Pvp Actin*/*Pea Actin* in the *Pseudomonas*-treated peas relative to the control (LB-treated pea) was plotted. Plots show data from one of the three independent biological repetitions. T-test was used to compare the means for significant differences. **=significant at p-value of <0.01.

### Downy mildew biomass analysis supports drenching data

We further investigated whether drenching peas with *Pseudomonas* reduced the total DNA of the pathogen. To assess this, the DNA of the *Pvp-Actin* gene, which plays a critical role in the pathogen’s structure, movement, and virulence, was quantified via qPCR. Consistent with the earlier spore count data, the *Pvp* DNA biomass analysis showed a significant (p < 0.05) decrease (95.7% less DNA compared to the control) in the *Pvp*-inoculated peas drenched with *Pseudomonas* compared to those drenched with LB medium (Fig. 7D).

### Foliar application of biocontrol agents or filtrates suppress downy mildew growth

In addition, we tested the direct effectiveness of foliar application of the biocontrol agents. Pea plants, which were infected with *Pvp* and expected to produce spores, were treated with either bacterial cells or their filtrates. The application of both cells and filtrates from all Bacillus strains reduced sporulation by 91 to 96.1% for cells and 85 to 89.7% for filtrates compared to the *Pvp*-infected control plants, which are not treated with *Bacillus* filtrates or cells. Similarly, *Pseudomonas* cells and filtrates reduced sporulation by 98.2% and 87.1%, respectively, significantly inhibiting *Pvp* sporulation in three separate trials (Figs. 8A-D and 9A-B, Supplemental Figs. S2 & S3).

**Figure 8:**
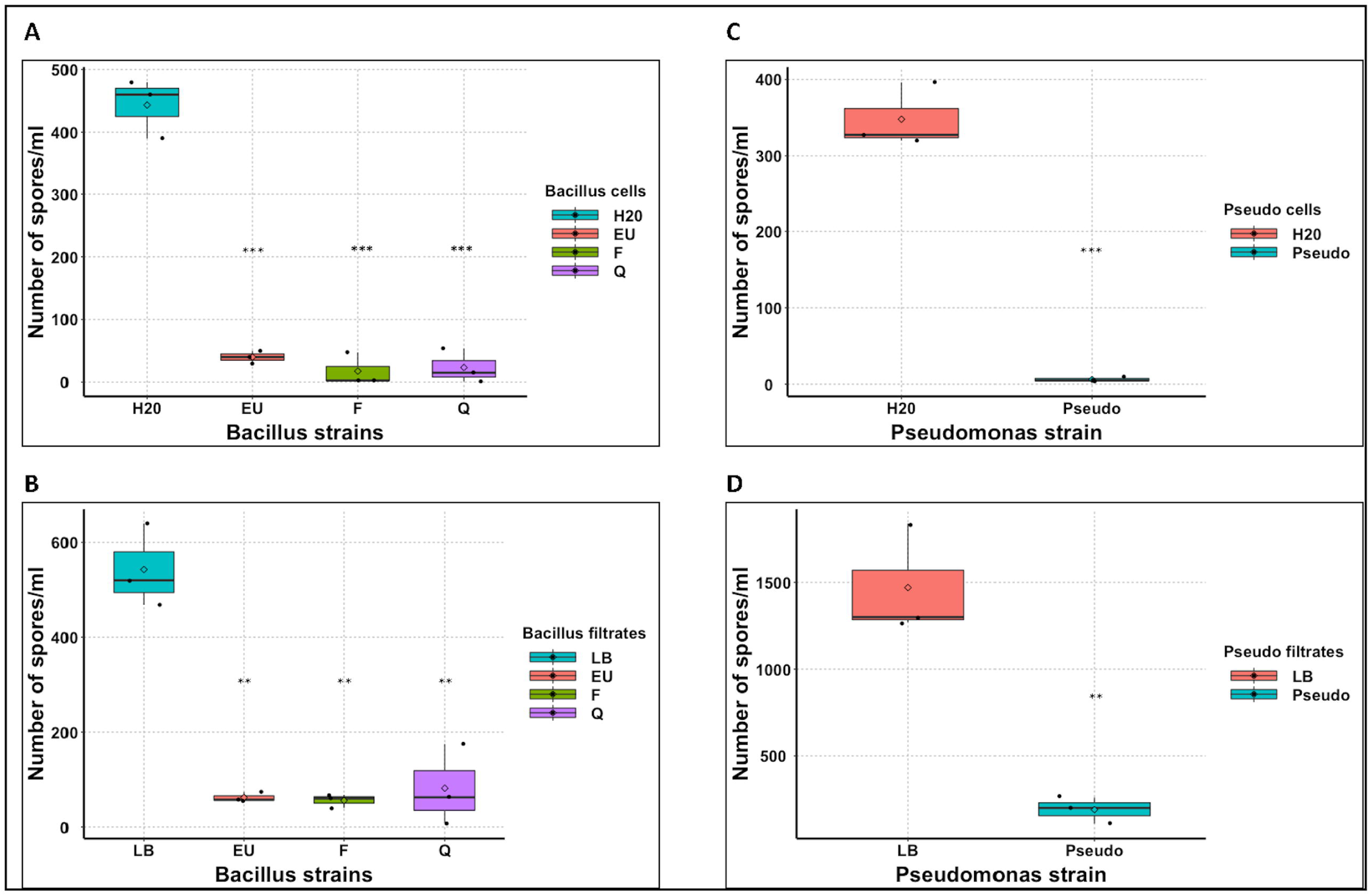
**Antagonism assay of foliar application of tropical *Bacillus* and *Pseudomonas* cells/filtrates on *Pvp*- inoculated pea plants**. 4-day old seedlings were inoculated with *Pvp* spores and grown in the growth cabinet. After 10 days, plants were sprayed with the biocontrol or the control and covered for 2 days to induce *Pvp* sporulation. After sporulation, spores were harvested and counted. **A**: Mean spore counts for plants sprayed with *Bacillus* cells and the control (H_2_0), **B**: Mean spore counts for plants sprayed with *Bacillus* filtrates and the control (LB), **C:** Mean spore counts for plants sprayed with *Pseudomonas* cells and the control, **D**: Mean spore counts for plants sprayed with *Pseudomonas* filtrates and the control. Plots show data from one of the three independent biological repetitions. T-test was used to compare the means for significant differences. **=significant at p-value of <0.01. ***=significant at p-value of <0.001.

**Figure 9:**
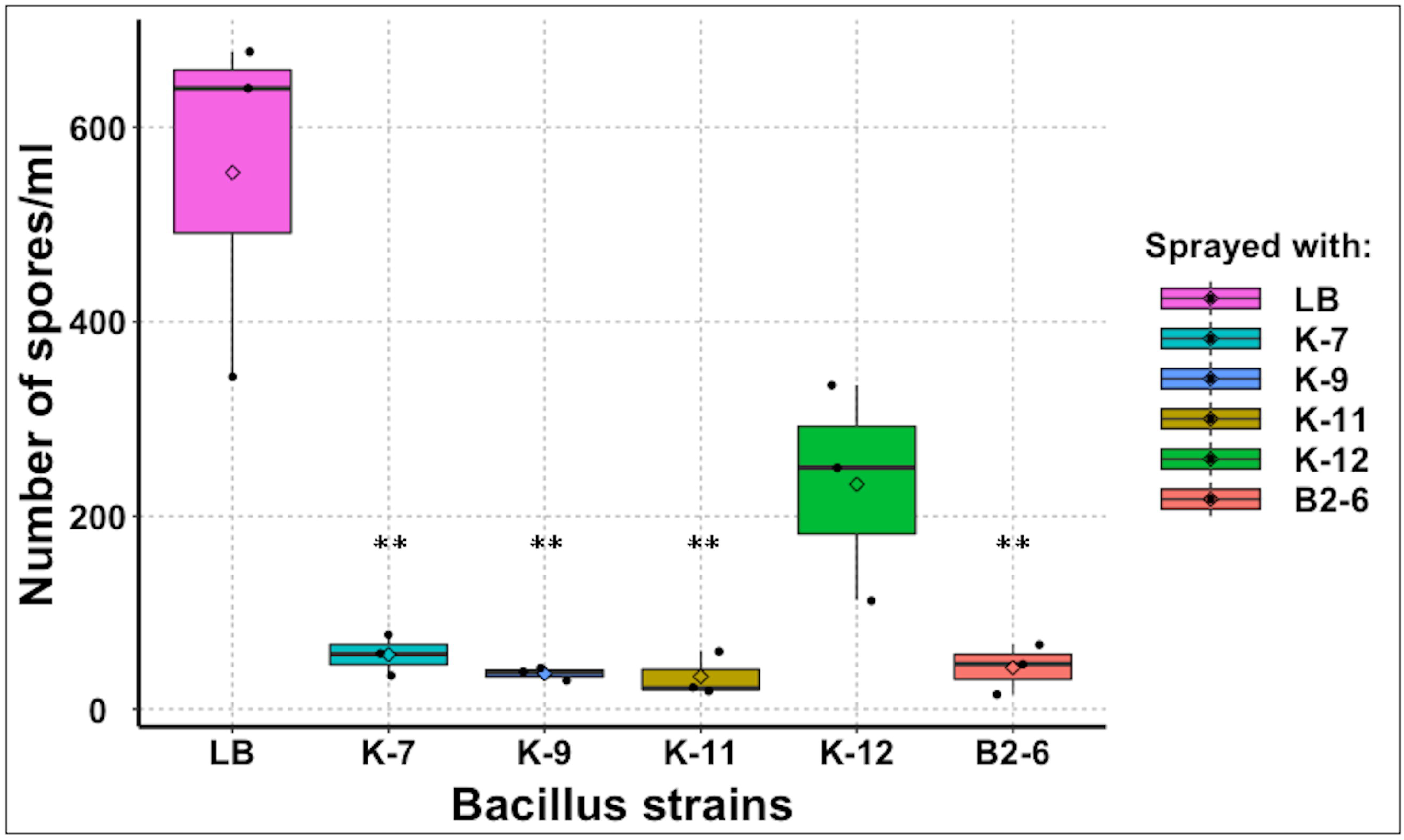
**Antagonism assay of foliar application of cold-loving *Bacillus* filtrates on *Pvp*- inoculated pea plants**. 4-day old seedlings were inoculated with *Pvp* spores and grown in the growth cabinet. After 10 days, plants were sprayed with the biocontrol or the control and covered for 2 days to induce *Pvp* sporulation. After sporulation, spores were harvested and counted. Mean spore counts for plants sprayed with filtrates of *Bacillus* strains (K-7, K-9, K-11, K-12 & B2-6) and the control (H20) are displayed in box plots. Plots show data from one of the three independent biological repetitions. T-test was used to compare the means for significant differences. **=significant at p-value of <0.01.

We also monitored the durability of the biocontrol agents by allowing the sporulated plants to grow for an additional 5 days in the growth cabinet. Remarkably, the pathogen did not recover on the biocontrol-treated plants, while the control plants retained *Pvp* spores (Figs. 10A-B).

**Figure 10:**
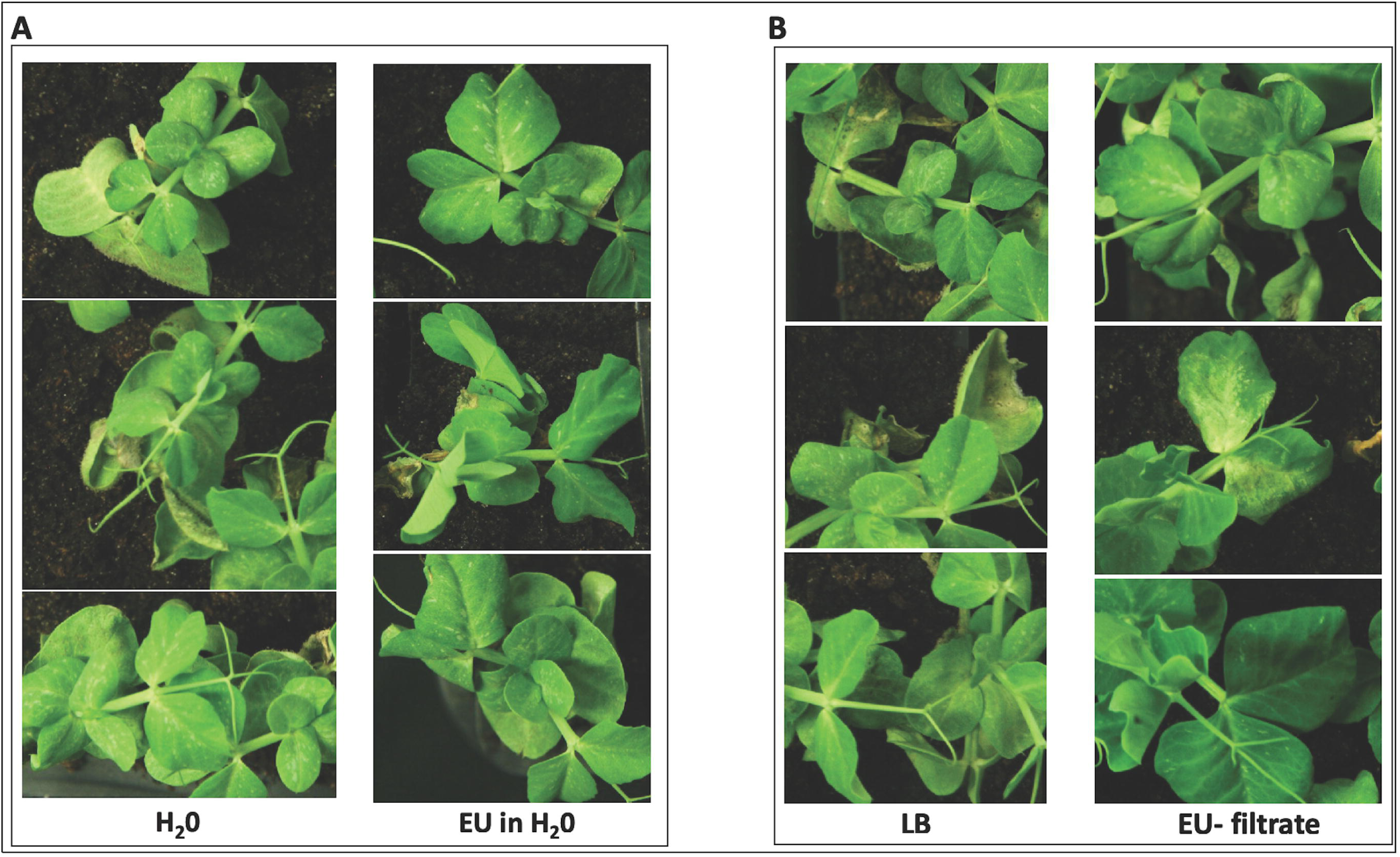
**Magnified images showing durability of EU07 antagonism on *Pvp* sporulation in pea plants**. 4-day old seedlings were inoculated with *Pvp* spores and grown in the growth cabinet. After 10 days, plants were sprayed with EU07 cells or filtrates. Control plants were sprayed with H_2_0 or LB. The plants were covered for 2 days to induce *Pvp* sporulation. After sporulation, the plants were uncovered and returned to the growth cabinet for 5 days. Images were taken after 5 days for those sprayed with EU07 cells or H_2_0 (**A**) and those sprayed with EU07 filtrate or LB (**B**).

### Dual applications of *Pseudomonas* and *Bacillus* strains demonstrate synergistic effect in downy mildew suppression

We conducted a study to determine if using both *Pseudomonas* and *Bacillus* bacterial strains together would have a greater impact on reducing pathogen growth compared to using them individually. We tested thisby using the filtered byproducts of these bacteria. Considering their optimal growth temperature, we combined tropical *Bacillus* and *Pseudomonas* strains, both of which have an optimal growth temperature of 28°C. The combined application of both bacterial strains significantly decreased the pathogen *Pvp* spore load by 88.3 to 97.3% compared to a control group using LB (Fig. 11). A synergistic effect was observed with the combined application showing a 27.6 to 46.7% greater reduction compared to when the bacteria were applied individually (Fig. 12)

**Figure 11:**
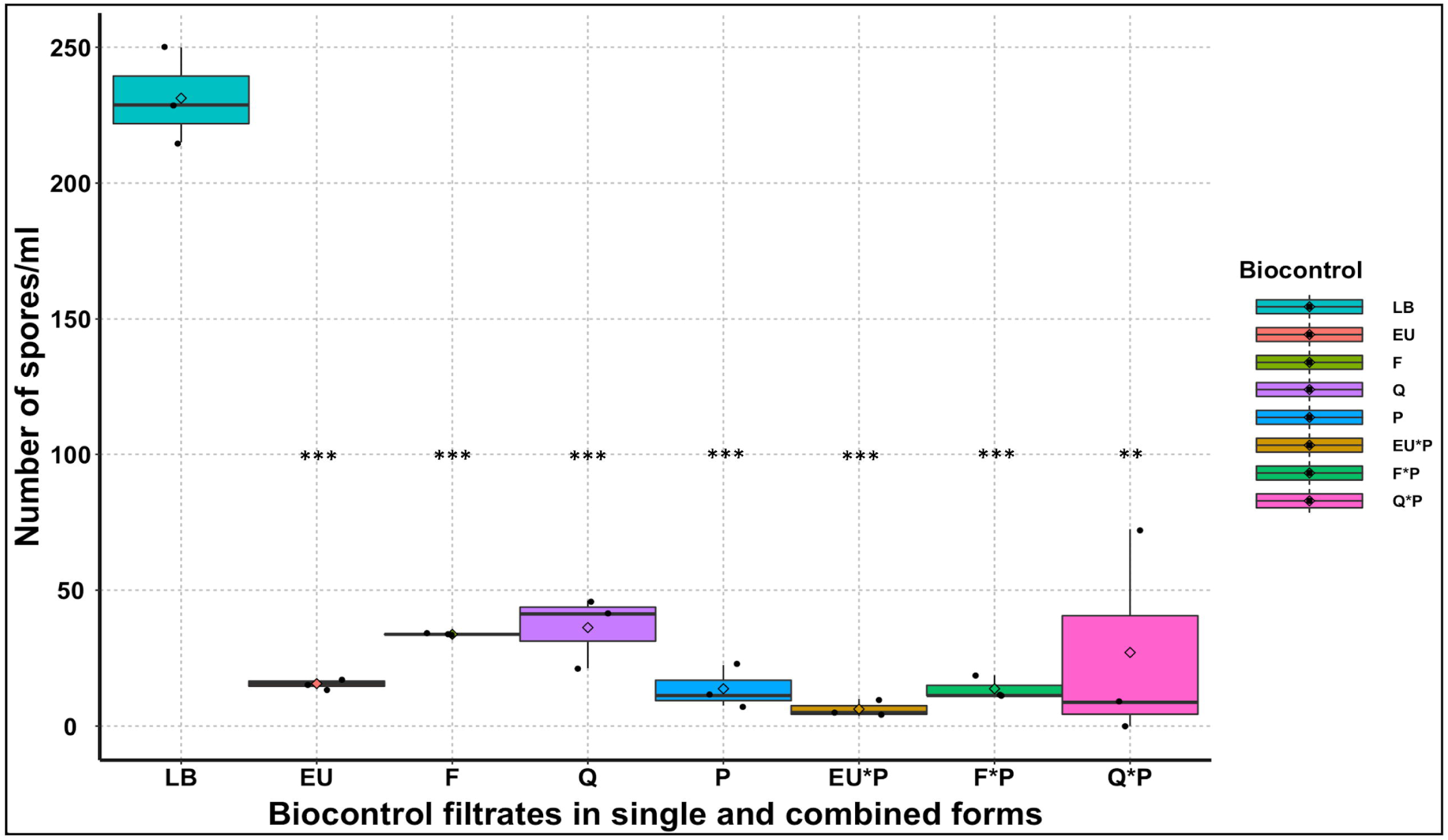
**Antagonistic effects of cocktail foliar application of *Bacillus* and *Pseudomonas* filtrates on *Pvp*- inoculated pea plants**. 4-day old seedlings were inoculated with *Pvp* spores and grown in the growth cabinet. After 10 days, plants were sprayed with the biocontrol agents in single or combined forms along with the control (LB). Plants were covered for 2 days to induce *Pvp* sporulation. Mean spore counts from plants sprayed with *Bacillus* (EU, F, Q) and/or *Pseudomonas* (P) and controls are displayed in boxplots. Plots show data from one of the three independent biological repetitions. T-test was used to compare the means for significant differences. **=significant at p-value of <0.01. ***=significant at p-value of <0.001.

**Figure 12:**
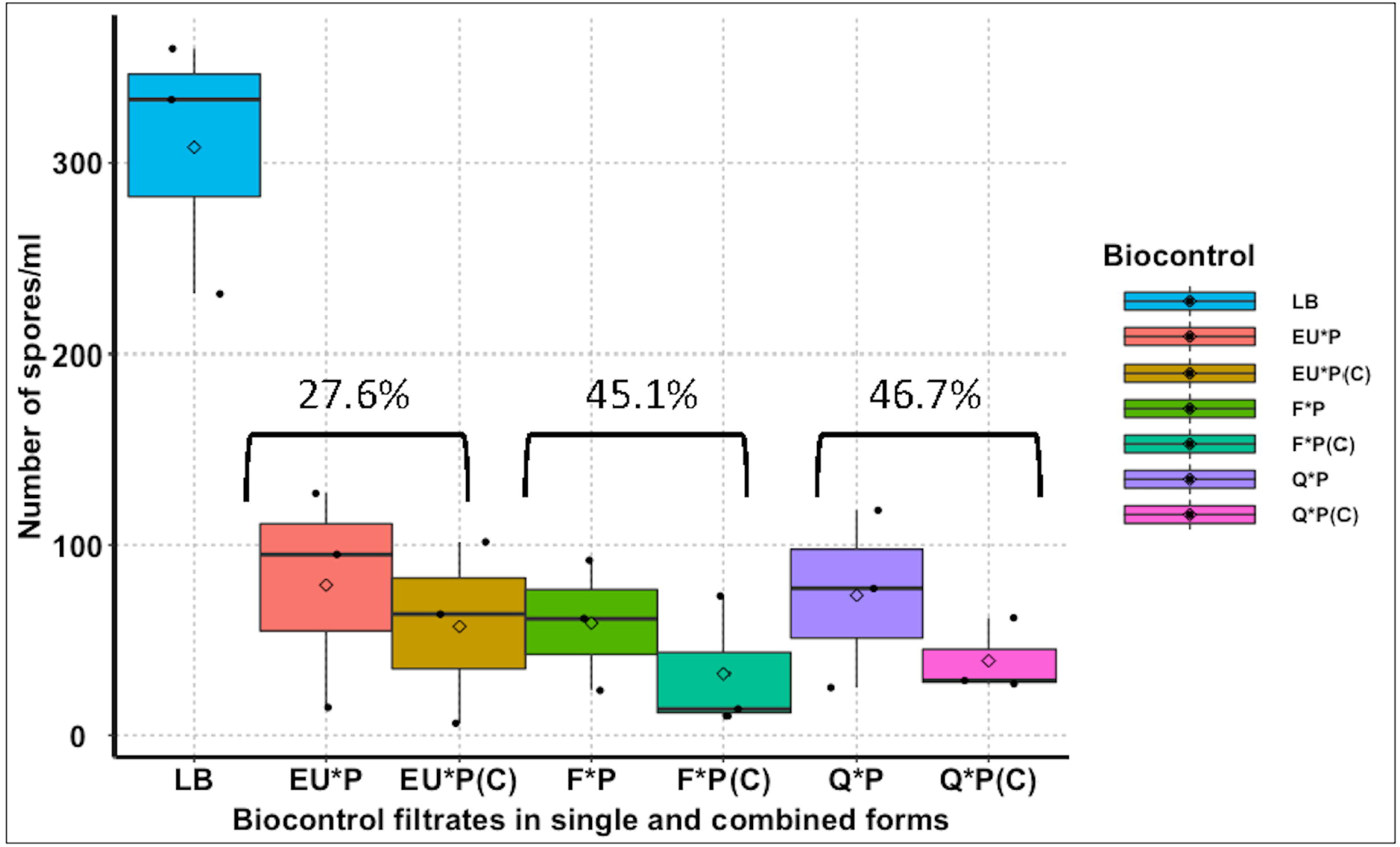
**Synergistic effects of combined foliar application of *Bacillus* and *Pseudomonas* filtrates on *Pvp*-inoculated pea plants**. Synergistic effects were calculated from the dual foliar application of *Bacillus* and *Pseudomonas*. Combinations with ‘C’ show those that were combined and sprayed as cocktails, while combinations without ‘C’ show those that were applied individually, and their mean effects calculated. Mean values from three independent biological repetitions were used to construct the box plots.

### Application of the biocontrol has no side effects on healthy pea

The *Bacillus* EU07 strain (both cells and filtrates) was tested for potential visual side effects after foliar applications on healthy pea plants. As shown in Fig. 13, no visual side effects were observed in the pea plants treated with either EU07 bacterial cells or corresponding filtrates compared to their respective controls. In fact, the biocontrol-treated plants, including the controls (mock), appeared as healthy and stress-free as the non-treated ones.

**Figure 13:**
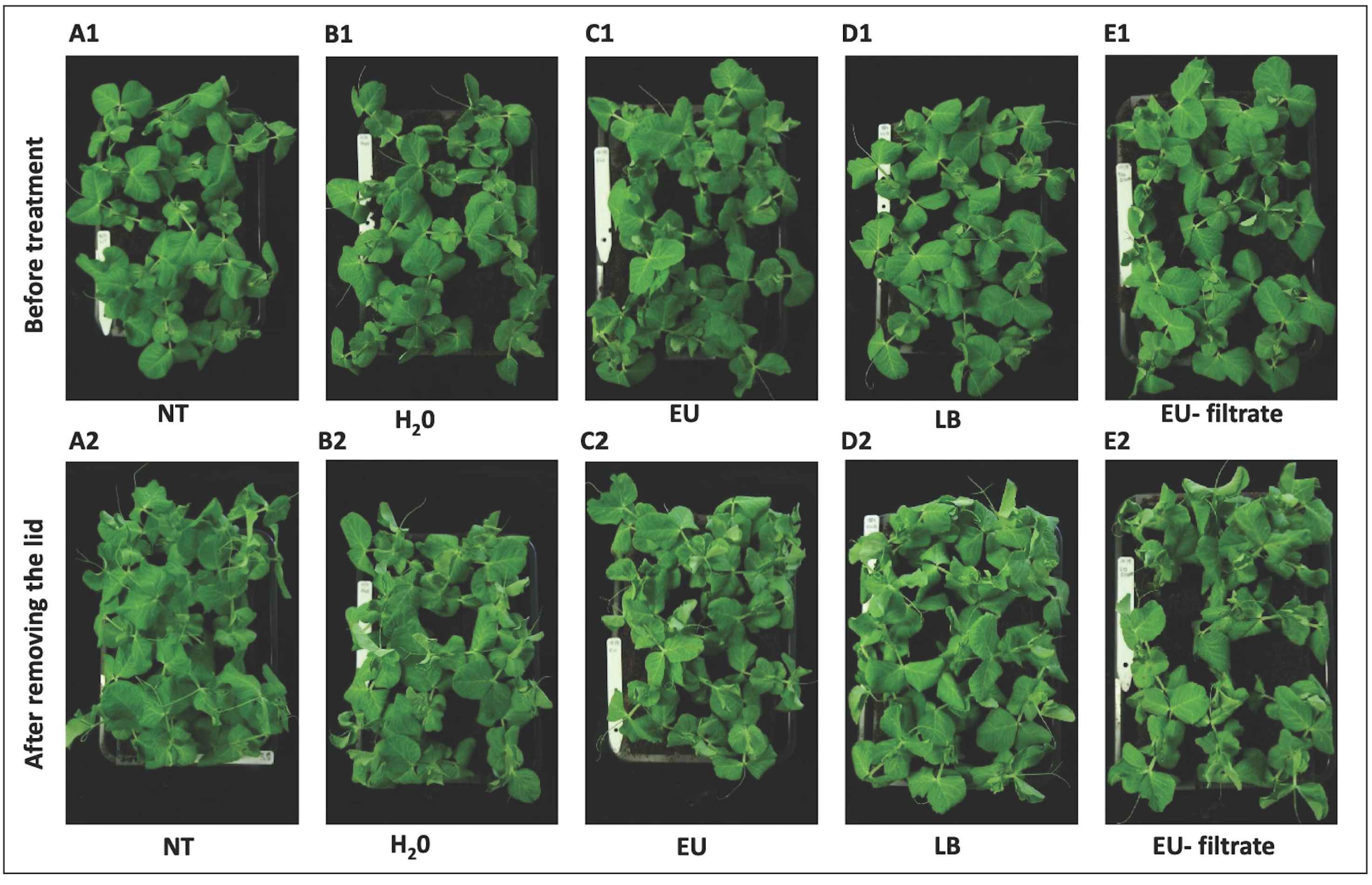
**Evaluation of negative effects of biocontrol sprays on healthy pea plants**. 4-day old seedlings were sown in pots and grown in the growth cabinet. After 10 days, pea plants were sprayed with the biocontrol. Plants were covered and moved to the growth cabinet for 2 days. Upper images show plants before being covered. Pea plants with no spray (**A1**), sprayed with H_2_0 (**B1**), EU07 cells (**C1**), LB (**D1**) and EU07 filtrates (**E1**). Lower images show pea plants after being covered for 2 days. Pea plants with no spray (**A2**), sprayed with H_2_0 (**B2**), EU07 cells (**C2**), LB (**D2**) and EU07 filtrates (**E2**).

### Genomic analysis confirms *P. fluorescens* and identifies the cold-adapted *Bacillus* K11 as *B. velezensis*

We used genome sequences to confirm the identity of the commercially purchased *P. fluorescens* LZB 065. Our genome assembly for LZB 065 was almost identical (with Average Nucleotide Identity (ANI) of 99.9948 %) to the genome of *P. fluorescens* type strain DSM 50090 (GenBank: GCA_007858165.1). Although the vendor provides no information about the provenance of LZB 065, it is therefore likely that it is derived from this type strain.

The genome sequence data for the *Bacillus* strain K11 provided confirmation that it belongs to the species *B. velezensis*. The TYGS webserver identified K11 as belonging to this species and it shares 97.9673% ANI with the type strain. Strain K11 is phylogenetically distinct from strains FZB32 and from EU07 and QST713 (Fig. 14). The most closely related genome sequence currently available is that of strain DE0372 (99.3861 % ANI), isolated from an environmental sample in North Carolina, USA, in 2018 (BioSample: SAMN11792532).

**Figure 14.**
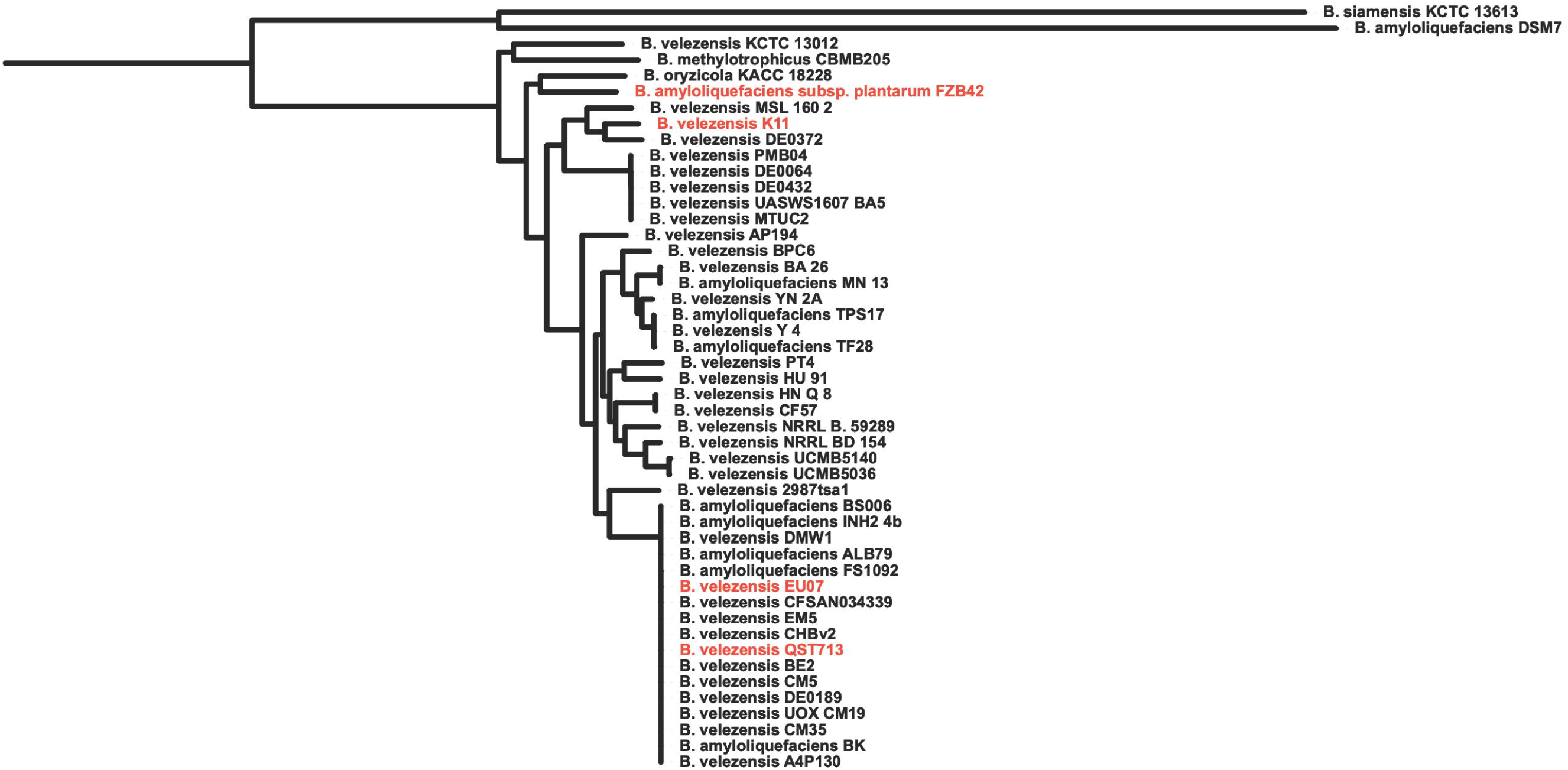
Phylogenetic tree *of Bacillus velezensis* strains, based on genome sequence data. The strains used in this study (FZB42, K11, EU07 and QST713) are highlighted in red. The tree was rooted by including *B. simanesis* and *B. amyloliquefaciens* type strains as an outgroup.

## Discussion

The use of microbial control biological agents (MBCA) is a safe and sustainable alternative to chemical pesticides. It not only protects crops against pathogens but also significantly reduces pollution and negative impacts of chemical pesticides on the environment (Jaiswal et al., 2022; Lahlali et al., 2022). Additionally, MBCA ensure the production of healthy and safe foods for human and animal consumption and well-being (Bale et al., 2008; Garvey, 2022). Current research is focused on exploring the untapped potential of MBCA (De Simone et al., 2021; El-Saadony et al., 2022; Lahlali et al., 2022). Bacteria and fungi such as *Bacillus*, *Pseudomonas*, *Streptomyces* and fungi such as *Trichoderma*, *Rhizophagus*, and *Clonostachys* have been tested and commercialized as biopesticides and bioprotectants against a wide range of plant pathogens (El-Saadony et al., 2022; Jangir et al., 2021; Thambugala et al., 2020).

No specific MBCA has been reported to be effective against the downy mildew pathogen in pulses, including pea crops. To address the question of whether MBCA can suppress this pathogen, we tested the potential of three strains of *Bacillus velezensis* and a strain of *Pseudomonas fluorescens* as biopesticides against the pea downy mildew pathogen *Peronospora viciae* f. sp. *pisi* (*Pvp*).

The antagonistic abilities of three *Bacillus* strains (EU07, FZB24, and QST713) and *Pseudomonas* to inhibit *Pvp* spore germination were demonstrated. The microbes were mixed with the *Pvp* spores and the mixtures were then incubated to assess the impact of the biocontrol on the spore germination percentage. This method was used because *Pvp* is an obligate pathogen and cannot be propagated without the host, making traditional *in vitro* bioassays using agar media unsuitable. However, the system used for the bioassays in this research has been employed previously by other researchers such as Bilir et al. (2019) and Telli et al. (2020). In these bioassays, the cells (pellets suspended in water) and filtrates (supernatant after centrifugation) of the *Bacillus* and *Pseudomonas* strains were separately tested, as the filtrates could contain antimicrobial metabolites. Interestingly, the cells and filtrates of all the potential MBCA showed complete inhibition of *Pvp* spore germination even at 50% concentration.

The positive antagonistic effects observed, especially with the filtrates, align with a significant body of literature explaining that the primary mechanism of direct antagonism of these microbial biocontrol agents is their natural ability to produce and use various antimicrobial substances such as lipopeptide, subtilin, bacilysin, mycobacillin, bacillomycin, fengycin, surfactin, and iturin to inhibit the growth and proliferation of pathogenic microorganisms (Hashem et al., 2019; Ntushelo et al., 2019; Shoda, 2000).

While *in-vitro* antagonism on *Pvp* has not been reported in the literature, the effectiveness of *Bacillus* and *Pseudomonas* spp and their filtrates has been demonstrated using agar-based in-vitro systems to be effective against various pathogens. For example, the application of *Bacillus* species significantly inhibited *Fusarium graminearum* by up to 79% (Jimenez-Quiros et al., 2022), *Botrytis cinerea* by up to 87% (Chen et al., 2019), and *Sclerotium rolfsii* by ∼88% with cells and 100% with filtrates (Sultana & Hossain, 2022).

Our *in vitro* assays with *Bacillus* and *Pseudomonas* strains demonstrated suppression of *Pvp* spore germination. However, this effect may vary in the plant-microbe interaction environment. Therefore, the antagonistic activities of the *Bacillus* and *Pseudomonas* strains against *Pvp* were further studied in the host crop, pea. The biocontrol applications were either by drenching *Pvp*-inoculated pea seedlings (before infection developed) with *Bacillus/Pseudomonas* broths or by foliar spraying their cells/filtrates on the inoculated plants (after infection developed). Drenching the soil with the MBCA was only significantly effective for cold-loving Bacillus K11 and P. fluorescens (approximately 90% reduction in spore load compared to the control). However, significant suppressions of *Pvp* sporulation in pea plants sprayed with all the strains of *Bacillus/Pseudomonas* (∼90%) or their filtrates (more than 80%).

The positive *in-planta* antagonism supports several studies that have shown that rhizobacterial *Bacillus* and *Pseudomonas* species can suppress a wide range of plant pathogens (Dragana et al., 2017; Gao et al., 2012; Mnif & Ghribi, 2015). For example, Núñez-Palenius et al. (2022) reported that foliar application of *B. subtilis* effectively controlled downy mildew disease in cucumber (caused by *Pseudoperonospora cubensis*) in a controlled environment. Kremmydas et al. (2013) also indicated in their research that *Pseudomonas fluorescens* strain X was able to suppress cucumber and sugar beet damping-off caused by the oomycete pathogen *Pythium ultimum*. The consistent results of *in vitro* and *in planta* antagonism assays in this research, in which both the cells and filtrates significantly suppressed *Pvp* growth and proliferation, suggest that one of the modes of action of these biocontrol agents could be their abilities to produce antimicrobial substances, as observed with their filtrates (Biniarz et al., 2018; Raaijmakers et al., 2010; Shafi et al., 2017).

Deravel et al. (2014) noted that two antimicrobial compounds, mycosubtilin and surfactin, obtained from the filtrates of two *B. subtilis* strains, were highly effective in controlling lettuce downy mildew disease caused by *Bremia lactucae*. Similar results were found in a study by Li et al. (2019), where surfactin and fengycin purified from another *Bacillus* strain were effective against grape downy mildew. Apart from the antibiosis mode of interaction, biocontrol agents can also use different antagonistic mechanisms, such as competing for space and nutrients, mycoparasitism, or indirectly priming/activating the host resistance genes, either separately or synergistically, to inhibit the growth and activities of pathogens (Bonaterra et al., 2022; Kohl et al., 2019; Legein et al., 2020; Roca-Couso et al., 2021).

In this study, we also assessed the persistence of antagonistic actions of biocontrol agents. The findings showed that *Pvp* did not visually recover on plants sprayed with biocontrol agents at 5 dpi, while the control plants still had *Pvp* spores on them. Bardin et al. (2015) stressed the need for further research on the durability of biocontrol agents to minimize potential failure or variations in their effectiveness, particularly in new environments. The positive results highlight the significant untapped potential that biocontrol agents offer in sustainable agriculture. In addition to investigating their durability, combining different biocontrol agents that share similar growth conditions as cocktails has been found to have synergistic effects, resulting in more effective antagonistic behaviour than when applied individually (Bardin et al., 2015; Xu et al., 2011). This is because each biocontrol agent exhibits unique features in how they demonstrate antagonistic activities; for example, some may produce distinctive types or quantities of antimicrobial substances or employ different combinations of antagonistic mechanisms. Combining them would harness all their individual attributes and positive interactive activities, leading to a more robust and efficient pesticidal effect on target pathogens (Kohl et al., 2019).

Synergistic effects of combining filtrates of tropical *Bacillus* strains (EU07, FZB24, and QST713) with *Pseudomonas* that have common peak growth temperature as foliar sprays on *Pvp*-inoculated peas were examined. The cocktail application significantly decreased *Pvp* spore load compared to the control and mixed application of the two biocontrol agents showed synergistic effects (27.7 to 46.7 % compared to individual application). Similarly, Abeysinghe (2009) indicated that cocktail application of *B. subtilis* with *P. fluorescens* strains showed higher plant protection against *Rhizoctonia solani* and *Sclerotium rolfsii* in *Capsicum annuum* (red pepper) than in the plants treated with either of the biocontrol agents alone (up to 45 %). Other researchers also reported increased antagonistic actions following application of combined biocontrol agents against different plant pathogens (Diaz- Manzano et al., 2022; El-Sharkawy et al., 2022; Palazzini et al., 2022; Panchalingam et al., 2022). Assemblage and use of different diverse biocontrol agents as consortia is an effective way to increase the efficiency and durability of microbial biocontrol agents (Sarma et al., 2015). However, compatibility and possible interaction of the proposed biocontrol agents to be combined needs to be studied to ensure there are no negative interactions from their combination that would result in reduced efficacy relative to their individual efficacies (Niu et al., 2020; Sarma et al., 2015).

Although biocontrol agents are widely considered safe and have little to no negative effects on the environment and ecosystems (Bhat et al., 2023; El-Saadony et al., 2022; Li et al., 2022), some researchers caution that since these microbes or their by-products are intentionally applied, often in high amounts, their biosafety, especially on non-target organisms, should be tested (Barat, 2011; Delfosse, 2005; Kiss, 2004; Winding et al., 2004). Therefore, in the present study, a simple biosafety analysis of the biocontrol agents (using EU07 *Bacillus* strain as a representative) was conducted. Following spraying of EU07 and its filtrate on healthy pea plants, no negative effects on the plants were observed compared to the control plants. This indicates that the type and dosage of the biocontrol agents used in this study are safe for use in crop protection, as also indicated by other researchers (Brutscher et al., 2022; Deravel et al., 2014; Lefevre et al., 2017).

## Conclusion

The lack of information on the efficacy of potential biopesticides and the lack of credible alternatives to chemicals for controlling downy mildew pathogens in pulses led to this research. We studied the effectiveness of various strains of *Bacillus velezensis* and *Pseudomonas fluorescens* in combatting pea downy mildew caused by *Pvp*. In laboratory tests, all the *Bacillus and Pseudomonas* strains and their filtrates completely inhibited *Pvp* spore germination. Further research involved treating *Pvp*-inoculated pea seedlings with biocontrol broth via soil drenching, which was found to be significantly effective only for cold loving *Bacillus* K11 and *P. fluorescens*, as indicated by spore assays and molecular biomass quantification. When the biocontrol agents were applied as foliar sprays on *Pvp*-inoculated pea plants, those treated with *Bacillus* strains, *P. fluorescens* or their filtrates showed a significant decrease in spore numbers compared to the control. Additionally, combining *Bacillus* strains and *P. fluorescens* resulted in a synergistic reduction of *Pvp* spore load. We also assessed the safety of using these biocontrol agents as biopesticides on healthy pea plants, and found no obvious negative effects, confirming their safety and environmental compatibility. This research, being the first on the biocontrol of pea downy mildew, will provide a crucial foundation for further studies. Importantly, cocktails of *Bacillus* strains and *P. fluorescens* could be effective immediately in controlling pea downy mildew disease, thus bolstering the health of a significant nitrogen-fixing crop in rotations.

## Acknowledgements

We would like to express our gratitude to Dr. Jane Thomas for collaborating closely with us in securing funding and at the outset of the project.

## Author contributions

MT conceived the research idea and designed the experiments with ECO, who conducted the majority of experiments and performed data analysis. CJQ assisted with experimental work, and ÖB revised and edited the manuscript. SK and BA isolated the cold-adapted bacterial strains, while AW provided bioinformatics support on pea downy mildew. TW, SA, and CD contributed to writing and refining the manuscript. DJS conducted bacterial genomics studies and managed submission of genomic data to public databases. CD and MT secured funding for the BBSRC LINK project (BB/T016043/1). All authors contributed to the writing and review of the manuscript and approved the final version for submission.

## Data availability statement

The data that support the findings of this study are available from the corresponding author on reasonable request. All genomic data are publicly available as described in the paper.

## Supplemental Materials

**Supplemental Figure S1**: **Magnified pictures from antagonism assay of drench- application of *Bacillus* broth on *Pvp*- inoculated pea plants**. 4-day old pea seedlings were inoculated with *Pvp* spores and sown in a standard compost. *Pseudomonas* broth or LB was applied immediately upon sowing the seedlings. After 10 days, plants were covered for 2 days to induce *Pvp* sporulation. After sporulation, images were taken for plants drenched with LB (**A**) or *Pseudomonas* broth (**B**).

**Supplemental Figure S2**: **Magnified pictures from antagonism assay of foliar application of EU07 on *Pvp*-inoculated pea plants**. 4-day old seedlings were inoculated with *Pvp* spores and allowed to grow in the growth cabinet. After 10 days, plants were sprayed with EU07 or water as a control and covered for 2 days to induce *Pvp* sporulation. After sporulation, images were taken for plants sprayed with H_2_0 (**A**) or EU07 cells (pellets suspended in water) (**B**).

**Supplemental Figure S3**: **Magnified pictures from antagonism assay of foliar application of EU07 filtrate on *Pvp*- inoculated pea plants**. 4-day old seedlings were inoculated with *Pvp* spores and grown in the growth cabinet. After 10 days, plants were sprayed with EU07 filtrate or LB as a control and covered for 2 days to induce *Pvp* sporulation. After sporulation, images were taken for plants sprayed with LB (**A**) or EU07 filtrate (supernatant after centrifugation) (**B**).

**Supplementary Table S1:**
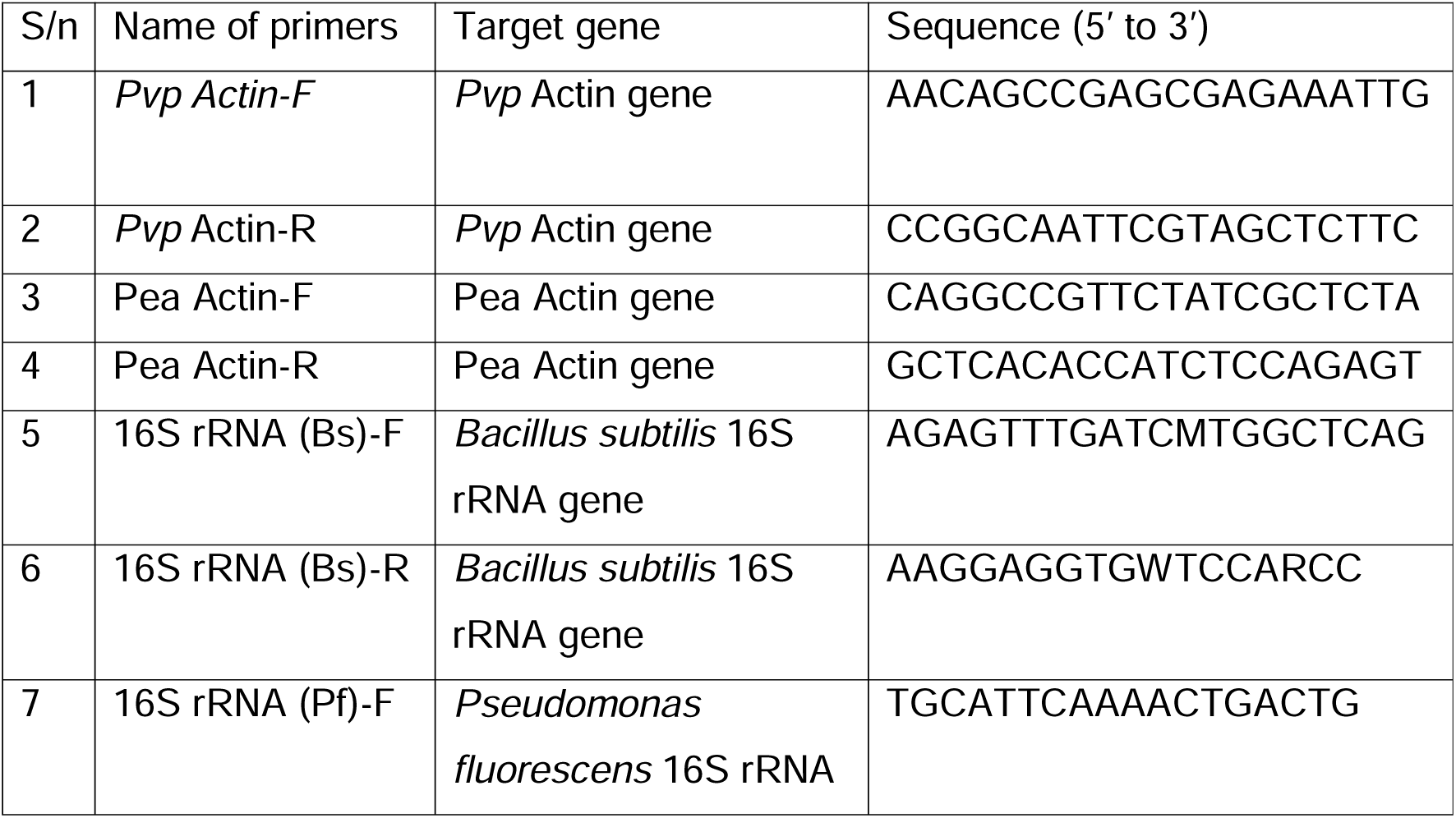
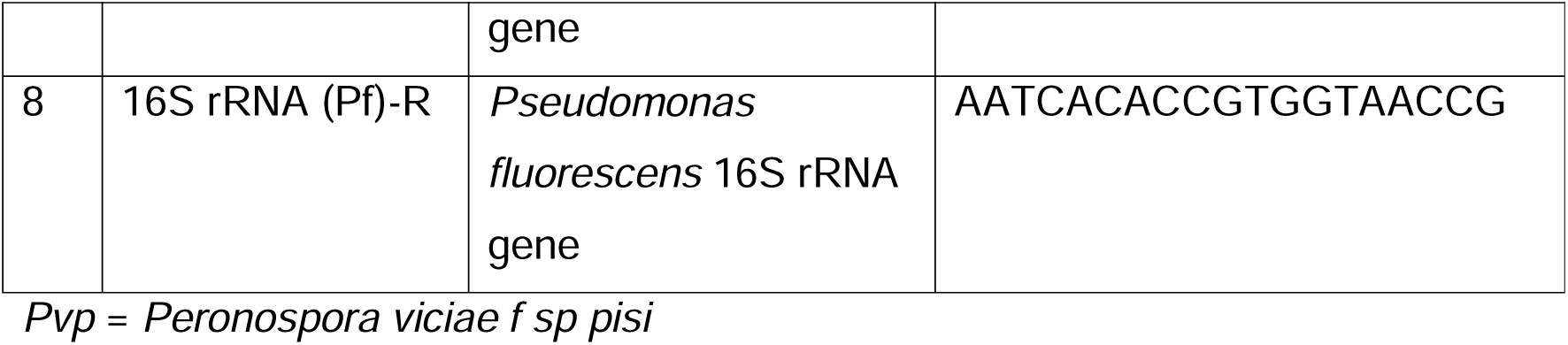
List of primers used in this research.

