## Supplementary figures and images for "Pea-Saving Partners: Bacillus and Pseudomonas combat downy mildew in pea crops"

### Suppl.Fig 1.jpg

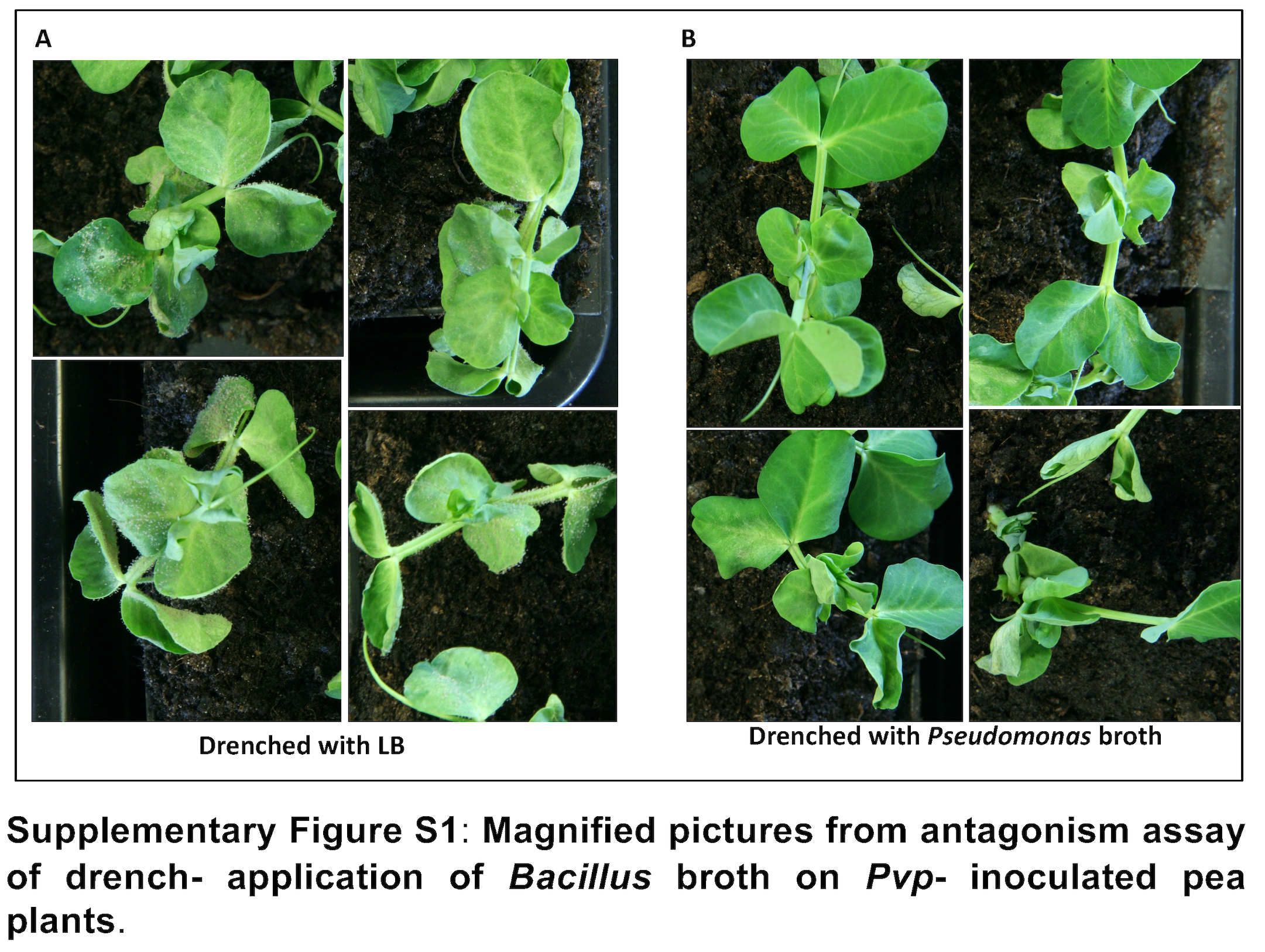

### Suppl.Fig 2.jpg

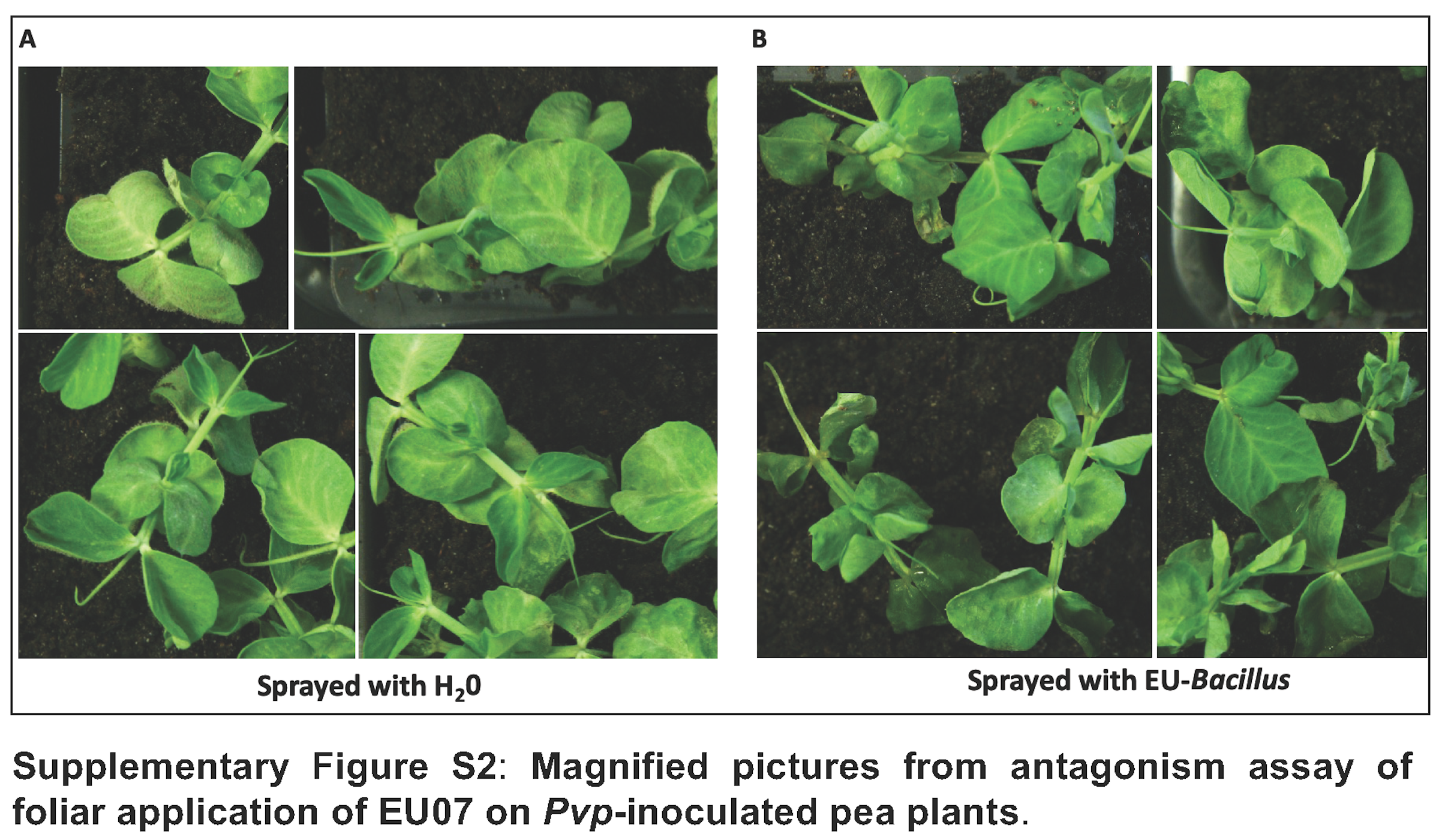

### Suppl.Fig 3.jpg

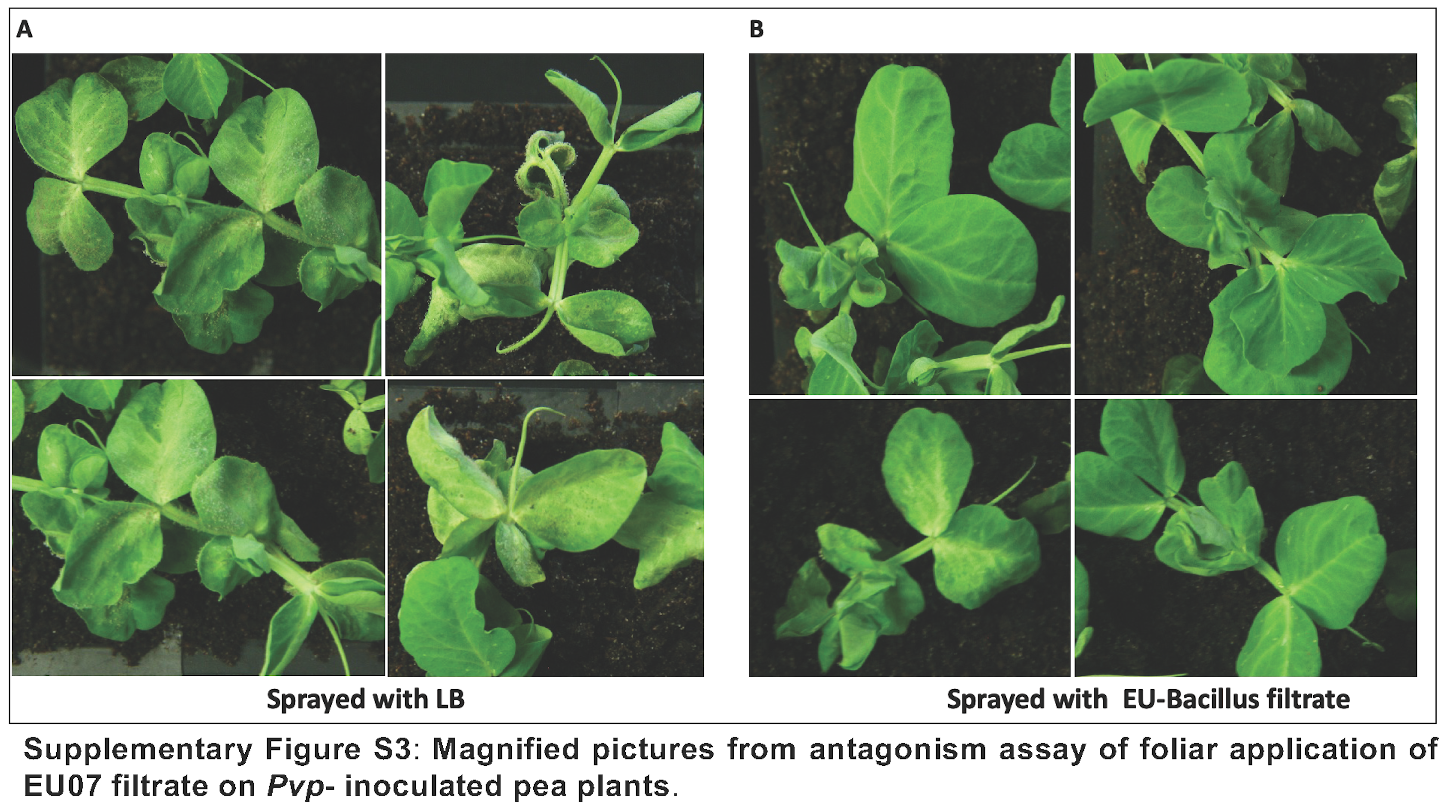
